# Inverted Assembly of the Lens Within Ocular Organoids Reveals Alternate Paths to Ocular Morphogenesis

**DOI:** 10.1101/2025.04.17.649366

**Authors:** Elin Stahl, Miguel Angel Delgado-Toscano, Ishwariya Saravanan, Anastasija Paneva, Joachim Wittbrodt, Lucie Zilova

## Abstract

The eye is a complex organ composed of two main structures – the retina and the lens. It forms by the invagination of the lens forming head surface ectoderm embedding into the forming optic cup. This “outside-in” mode of morphogenesis ensures that the light focusing lens is positioned centrally inside of the eye in the highly constrained environment of the developing embryo. Advances in stem cell biology in the last decade introduced organoids as model to study organogenesis under normal and diseased conditions. However, even though strikingly similar at some points, it remained elusive to which extend the generation of individual structural features in organoids recapitulates *in vivo* organogenesis. Here we describe the generation of fish ocular organoids composed of both, lens and retina, using pluripotent embryonic cells from medaka (*Oryzias latipes*). Formation of the organoid lens followed the key molecular features of the process *in vivo*, including the establishment of lens progenitor cells and their subsequent differentiation into lens fiber cells. In a process dependent on the coordinated activity of BMP and FGF signaling, lens formation in ocular organoids was marked by the expression of key genes implicated in organismal lens development. Despite adhering to the basic molecular machinery of lens formation *in vivo*, the morphogenesis into a spherical lens followed an “inside-out” mode. Lens progenitor cells were initially established and differentiated into a spherical lens directly inside of the retina. Subsequent displacement of the lens from the center of the organoid towards its surface ultimately led to the formation of a cup-like shaped retina with a centrally positioned lens. Our study highlights that the self-organization of the organoid can favor routes that were not selected for in the developing embryo. Those routes can lead to an alternative, though highly similar outcome with the respect to achieving specific structural features in an unconstrained, embryo-free environment.

## Introduction

The tissues of the embryo develop in an evolutionarily selected confined space with robust physical and biochemical constraints. Precise spatio-temporal gradients of growth factors aligned with complex mechanical forces guide cell and tissue interactions, permitting the coordinated formation of functional tissues and organs. Eye development is an example of complex organ formation relying on the coordinated interactions of tissues of different embryonic origin. This ultimately results in the positioning of a light-focusing lens in the center of the light-sensing retina (Cardozo et al., 2023; Casey et al., 2021; Chow and Lang, 2001; Cvekl and Ashery-Padan, 2014).

While the lens originates from the head surface ectoderm (SE), the retina originates from optic vesicle (OV), a lateral evagination of the forming diencephalon. In vertebrates, eye development is initiated at the end of gastrulation when the anterior ectoderm is patterned into the neural plate, non-neuronal ectoderm and the neural plate border at their interface. Lens-committed progenitors are first found in the anterior neural plate border, also called the anterior pre-placodal region (Bailey and Streit, 2005; Bhattacharyya et al., 2004; McCabe and Bronner-Fraser, 2009; Schlosser, 2006; Toro and Varga, 2007). Retina-committed progenitors, on the other hand, are established within the anterior neural plate, by the specification of the eye field, followed by the evagination of the optic vesicle (Adelmann, 1937; Eagleson et al., 1995; Kenyon et al., 2001; Li et al., 1997). Once the retina-forming OV contacts the lens-competent ectoderm, interactions of OV, SE and the surrounding tissues result in the formation of the lens placode within the head surface ectoderm. The concomitant invagination of lens placode and morphogenesis of the OV then lead to the formation of the lens vesicle in the center of the forming optic cup (Casey et al., 2021; Chow and Lang, 2001; Cvekl and Ashery-Padan, 2014; Gunhaga, 2011). Initially the lens vesicle is composed of lens progenitor cells. It is further patterned into lens epithelium at the anterior part of the lens, while posteriorly located progenitors differentiate into lens fiber cells (Kallifatidis et al., 2011; Zhou et al., 2006). The differentiation of lens fiber cells is characterized by the production and accumulation of crystallin proteins, the main structural proteins of the lens (Cvekl et al., 2015; Donaldson et al., 2017). This transition is marked by the extensive elongation of cells (Cheng et al., 2017; Fudge et al., 2011; Rao and Maddala, 2006) and the subsequent degradation of all organelles and de-nucleation (Bassnett, 2009; Brennan et al., 2021; Chaffee et al., 2014; Wride, 2011), minimizing light scattering and providing optimal refractive properties, all together resulting in the optical properties of the lens (Bassnett et al., 2011).

Several studies have shown that the conserved process of lens formation depends on the intricate interplay of intrinsic factors and extracellular signalling pathways (Cvekl and Ashery-Padan, 2014; Cvekl and Zhang, 2017; Lang, 2004; McAvoy et al., 1999; Ogino and Yasuda, 2000). The transcription factors *Pax6*, c-*Maf, Sox1, Prox1, Hsf4, Six3*, *Foxe3*, *Pitx3*, *Sox2* characterise early lens development and are directly involved in the processes of initial lens morphogenesis and lens fiber cell differentiation (Cvekl and Duncan, 2007; Cvekl and Zhang, 2017; Lang, 2004; Ogino et al., 2012; Ogino and Yasuda, 2000). Establishment of lens-competent progenitors as well as differentiation of lens fiber cells relies on the interplay of the FGF, BMP, and WNT signalling pathways (Faber et al., 2001, 2001; Furuta and Hogan, 1998; Grocott et al., 2011; Gunhaga, 2011; Jarrin et al., 2012; Kreslova et al., 2007; Le and Musil, 2001; Litsiou et al., 2005; Lovicu and McAvoy, 2005; McAvoy et al., 1999; Rajagopal et al., 2009).

Although many aspects of lens formation are well characterized, addressing the molecular mechanisms in the complex environment of the developing embryo is often challenging. Advances in stem cell and developmental biology allowed the introduction, fast development and prominent use of organoids as new models to study complex processes of *ex vivo* organ formation. Those allow to gain insight into cellular and molecular processes in normal and diseased conditions in a controlled and experimentally accessible *in vitro* environment (Chen et al., 2024; Hofer and Lutolf, 2021; Kretzschmar and Clevers, 2016; Sahu and Sharan, 2020; Zhao et al., 2022). Mouse, human and fish pluripotent embryonic stem cells have been shown to self-organise into retinal tissue when aggregated and cultured under minimal growth factor 3D suspension culture conditions. Retinal organoids recapitulate key aspects of embryonic development with respect to cell fate commitment, tissue patterning, cell differentiation and physiological functions (Eiraku et al., 2011; Kuwahara et al., 2015; Nakano et al., 2012; Völkner et al., 2016; Zilova et al., 2021). Since the culture conditions for the generation of retinal organoids favor neuroectodermal differentiation over non-neuronal ectodermal fates, they did so far not allow for concomitant lens formation (Eiraku et al., 2011; Kuwahara et al., 2015; Nakano et al., 2012; Zilova et al., 2021).

Generation of lens-like structures (lentoid bodies) by stepwise differentiation of human ES or iPS cells has been reported (Cvekl and Camerino, 2022; Dincer et al., 2013; Fu et al., 2017; Leung et al., 2013; Yang et al., 2010). However, most lens differentiation protocols directly favor the promotion of placodal fate on the expense of the neuronal fate (Dincer et al., 2013; Fu et al., 2017; Yang et al., 2010). This rendered the generation of more complex organoids composed of cell types of diverse embryonic origin challenging. Interestingly, two independent studies had reported the appearance of lens marker-expressing cells in close vicinity to retina tissue in 3D organoid culture (Gabriel et al., 2021; Mellough et al., 2015). However, considering the complex nature of tissue interactions required for the coordinated formation of both lens and retina *in vivo*, this raises the intriguing question of how a lens is formed in neuronal organoids and to what extend that process recapitulates *in vivo* organ formation.

Here we used organoids derived from *Oryzias latipes* blastula cells as a model system to investigate the formation of ocular organoids composed of both, retina and lens. By following the individual populations of lens- and retina-committed progenitors, we found that similar to the *in vivo* situation, lens and retina originate from spatially confined and separated populations of progenitor cells. The subsequent differentiation into their respective lineages was found to be dependent on BMP and FGF signaling pathways. Their differentiation and morphogenesis are characterized by the expression of key regulatory factors implicated in eye (lens and retina) organogenesis, however following an entirely different morphogenetic route towards forming an eye cup enclosing a centrally positioned lens.

## Results

### Fish retinal organoids form embedded lenses under specific culture conditions

We have previously reported the generation of retinal organoids from fish pluripotent embryonic cells grown in 3D suspension culture in minimal growth factor-containing media. Such organoids acquire anterior neural fate and consist of retina and presumptive forebrain tissue (Zilova et al., 2021). Here, we modified our previously described protocol (Figure 1a) in order to generate organoids, that differentiated into retinal and lens tissues.

**Figure 1.**
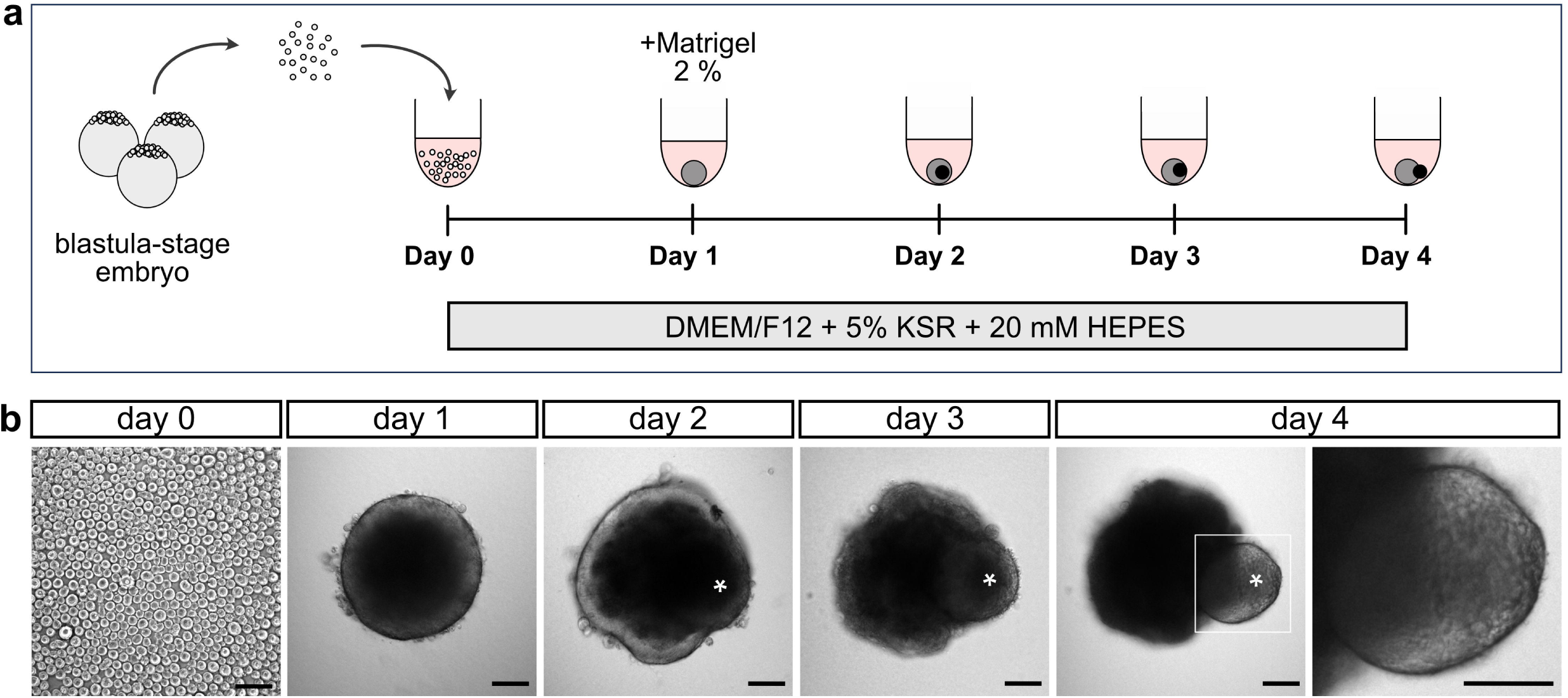
Generation of organoids from medaka pluripotent embryonic cells. **(a)** Schematic representation of organoid generation. At day 0, pluripotent embryonic cells were extracted from blastula-stage medaka embryos and seeded in U-bottom, low-binding 96-well plate into DMEM/F12 media supplemented with 5 % KSR and 20 mM HEPES pH 7.4. At day 1 aggregates were supplemented with 2 % Matrigel. **(b)** Bright-field pictures of indicated stages of organoid formation showing the appearance of lens-like structure (asterisk). KSR, knockout serum replacement. Pictures acquired with wide-field microscope (Mi8; Leica). Scale bar 100 μm.

Aggregation of fish primary pluripotent cells in DMEM/F12-based differentiation media supplemented with 5% knockout serum replacement and 20 mM HEPES pH 7.4 (see Materials and Methods for detailed composition) resulted in the efficient formation of retinal organoids containing a single transparent, spherical lens-like structure surrounded by neuronal tissue (Figure 1b). The lens-like structure was clearly visible at day 4 but first signs of its formation could be observed from day 2 onwards (Figure 1b, asterisk). We showed that organoids were composed of both retina and lens tissue, arranged in a way that was reminiscent of the tissue arrangement in the embryonic eye by using reporter lines and molecular markers. Differentiated lens cells expressing crystallin were labelled in the *Gaudí^RSG^* transgenic reporter line (Centanin et al., 2014) that carries a *crystallin::ECFP* (*Cry::ECFP*) reporter. Lens fiber cells were identified with the “lens fiber cell” marker antibody ZL-1 (referred to as LFC), and early and late lens cells were identified with an anti-Prox1 antibody (Figure 2a and 2b). Prox1 is expressed by cells of the lens placode and differentiating lens fiber cells of different species including mouse, zebrafish and medaka (Mikula Mrstakova and Kozmik, 2024; Pistocchi et al., 2008; Wigle et al., 1999). Anti-LFC labels lens fiber cells in zebrafish and is suggested to target one of the crystallin proteins (Greiling et al., 2010; Shi et al., 2006). In the medaka embryo, the expression of both, crystallin promoter-driven ECFP and LFC marker was observed exclusively in the lens 3 days post-fertilization (dpf) (Figure 2a). Prox1 immunolabeling was detected in the lens and a subpopulation of differentiating retinal cells in the central region of the retina (Figure 2a). The expression of all three lens markers was detected in day 4 ocular organoids (Figure 2b and 2c), demonstrating that the spherical structure formed by the organoids is indeed the lens. ECFP labeling was sparse (25 % *Cry::ECFP*, Figure 2b) reflecting the mode of generation of the organoids by a combination of 25 % cells carrying the *Cry::ECFP* reporter and 75 % wild-type cells (see Materials and Methods for details).

**Figure 2.**
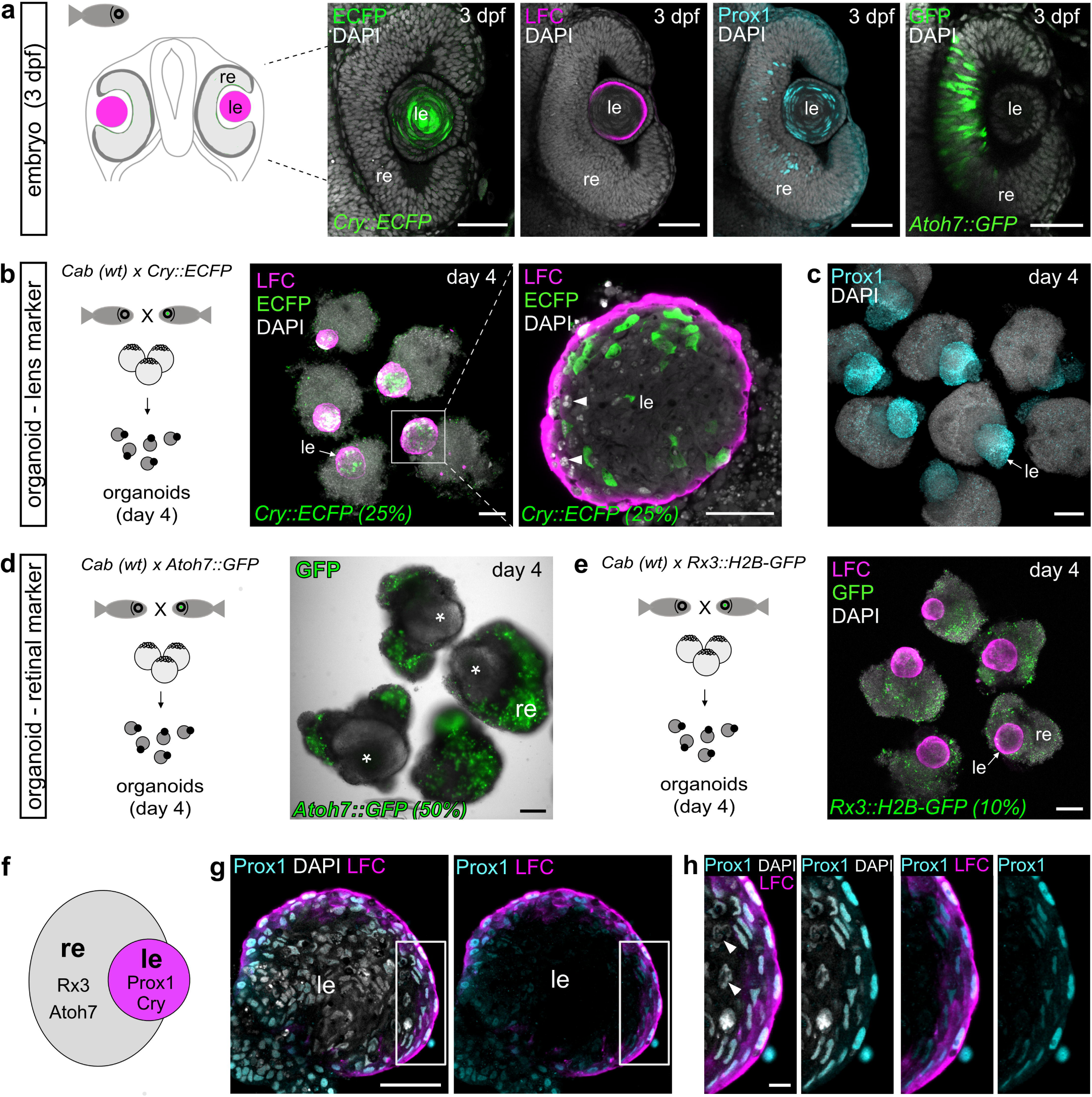
Fish retinal organoids form lenses. **(a)** Expression of lens-specific markers (*Cry::ECFP*, lens fiber cell (LFC) and Prox1) and retina-specific marker (*Atoh7::EGFP*) in medaka embryonic eyes at 3 dpf. Optical sections showing the expression of indicated markers detected by whole-mount immunohistochemistry with the use of anti-GFP (green), anti-LFC (magenta) and anti-Prox1(cyan) antibodies, co-labelled with DAPI nuclear stain (grey). **(b)** Organoid generation from crystallin reporter line (*Cry::ECFP*). Expression of lens-specific markers (*Cry::ECFP* and LFC) in day 4 organoids detected by anti-GFP and anti-LFC antibodies. Represented by maximal projection for overview and single optical section for the detail of the lens. **(c)** Expression of Prox1 in day 4 organoids derived from wild-type medaka cells detected with anti-Prox1 antibody, co-stained with DAPI nuclear stain; displayed as maximal z-projection. **(d**) Organoid generation from retina-specific reporter line (*Atoh7::EGFP*). Overlay of bright-field and wide-field images showing EGFP fluorescence in *Atoh7*-expressing cells, lens indicated with asterisk. **(e)** Organoids generated from retina-specific reporter line (*Rx3::H2B-GFP*) labeled with anti-GFP and anti-LFC antibody displayed as maximal z-projection. **(f)** Schematic representation of distribution of retina- and lens-specific markers in day 4 organoids. **(g)** Optical section of day 4 organoid lens labeled with anti-Prox1 and anti-LFC antibodies, co-labeled with DAPI, showing elongated Prox1^+^/LFC^+^ cells on the surface of the lens. **(h)** Enlargement of indicated area in g. For *Cry::ECFP*, *Atoh7::EGFP* and *Rx3::H2B-GFP*. The percentage (%) indicates how many cells carry the indicated transgene. Arrow in b, c and e indicates position of the lens. Arrowheads in b and h indicate abnormally looking nuclei. le, lens; re, retina; wt, wild-type; LFC, lens fiber cell. Scale bar 50 μm in a, g and enlargement of b, 100 μm in b-e, 10 μm in h.

The transgenic reporter lines *Atoh7::EGFP* (Del Bene et al., 2007) and *Rx3::H2B-GFP* (Rembold et al., 2006) were used to identify retinal tissue in lens carrying organoids. In embryos *Atoh7::EGFP* activity was confined to the retina, labelling differentiating retinal neurons at 3 dpf (Figure 2a). In organoids, labeled retinal cells surrounded the forming lens, reminiscent of the situation in the eye. To address the distribution of retinal and lens cells we used blastula cells from *Atoh7::EGFP* and *Rx3::H2B-GFP* retina-specific reporter lines (Figure 2d and 2e) and co-labeled *Rx3::H2B-GFP*-derived day 4 organoids with anti-LFC antibody (Figure 2e).

Analysis of the organoid lens revealed a great similarity with structural features of the developing embryonic lens, such as the elongation of Prox1^+^/LFC^+^ cells, typical for differentiating lens fiber cells and atypically looking nuclei indicating disorganization of nuclear material, a process tightly connected to lens fiber cell differentiation (Figure 2b and 2h, arrowhead).

In 462 organoids derived from 38 independent experiments we observed lens formation, in 83.5 % (n=386) of organoids. A careful evaluation of different culture conditions and their impact on the lens formation revealed that lens formation depends on supplementation of the DMEM/F12-based differentiation media with 20 mM HEPES buffer (Figure 2-figure supplement 1a and 1b). While organoids grown without HEPES supplementation never formed lenses (n=63 organoids in 6 independent experiments, see also Zilova et al., 2021), 92 % of organoids grown in media supplemented with HEPES (n=83 organoids in 6 independent experiments) formed lenses (Figure 2-figure supplement 1b).

We tested the impact of extracellular matrix (ECM) supplementation on lens formation, as existing protocols for generation of lentoid bodies from human pluripotent embryonic stem cells mostly rely on supplementation with ECM proteins (Dincer et al., 2013; Fu et al., 2017; Leung et al., 2013; Yang et al., 2010), usually in form of Matrigel. Interestingly, lens cell fate specification as well as lens fiber cell differentiation, judged by the expression of Prox1 and LFC, occurred independently of Matrigel (2 % final concentration) supplementation in medaka-derived organoids (Figure 2-figure supplement 1d and, f). Matrigel impacted mostly on the organization and thickness of Rx2^+^ retinal neuroepithelium localized on the surface of the organoids (Figure 2-figure supplement 1c). Measurement of the size of the lens (diameter of LFC^+^ lens sphere) at day 2 and day 5 showed that Matrigel supplementation did not impact the size of the lens (Figure 2-figure supplement 1d-e) or its overall organization analyzed by immunolabeling (Prox1, LFC, DAPI) at day 4 (Figure 2-figure supplement 1f).

### The lens originates from a spatially distinct population of placodal progenitors located in the central region of the organoid

We further investigated the origin of lens cells and how its formation compares to lens development *in vivo* in the developing organism. In the embryo, the lens originates from the lens competent head surface ectoderm (SE, magenta), which directly abuts the evaginating optic vesicle (OV, prospective retina, green) (Figure 3a). OV and SE invaginate in a coordinated way to form a lens sphere surrounded by the retina. In medaka embryos, those processes occur between 1 dpf and 2 dpf (Figure 3a, 3b). Concomitant with the completion of lens formation, the differentiation of lens fiber cells is initiated and marked by the expression of crystallin proteins (LFC; cyan) (Figure 3b). To track the distribution of lens and retinal progenitors, we generated organoids from the *Rx3::H2B-GFP* reporter line, which highlights retina-committed cells by nuclear GFP expression (Figure 3-figure supplement 1a and b). To visualize lens-committed progenitor cells within the SE (Mikula Mrstakova and Kozmik, 2024), we co-labeled those organoids with an anti-Prox1 antibody (Figure 3-figure supplement 1b) and followed the expression of *Rx3*-specific GFP and Prox1 from day 1 to day 2 (Figure 3c). Retinal progenitor cells were distinctly localized in the cortical region around the surface of day 1 organoids (Figure 3-figure supplement 1d, Figure 3c, Video S1), while lens progenitors (expressing Prox1) were not detectable at this stage (Figure 3c). Expression of retinal markers was confined exclusively to the cortex and was never detected in the central part of the organoid (Figure 2b, Figure 3-figure supplement 1d, Video S1). From day 1.5 onwards (stage S22 - 1 day and 14 h) (Iwamatsu, 2004), embryos as well as organoids (30 hours post aggregation) started expressing Prox1 (Figure 3b and 3c). In the embryo, Prox1 expression was localized to the surface of the head, labeling the presumptive lens region (Figure 3b). In the organoid, the expression was confined to the central core of the organoid with Prox1^+^ cells organized into a sphere (Figure 3c). Prox1 expression in the central core of the organoids was followed by the expression of lens fiber cell differentiation marker LFC at day 2, concomitant with the onset of Prox1 and crystallin expression in the embryo (Figure 3b and 3c).

**Figure 3.**
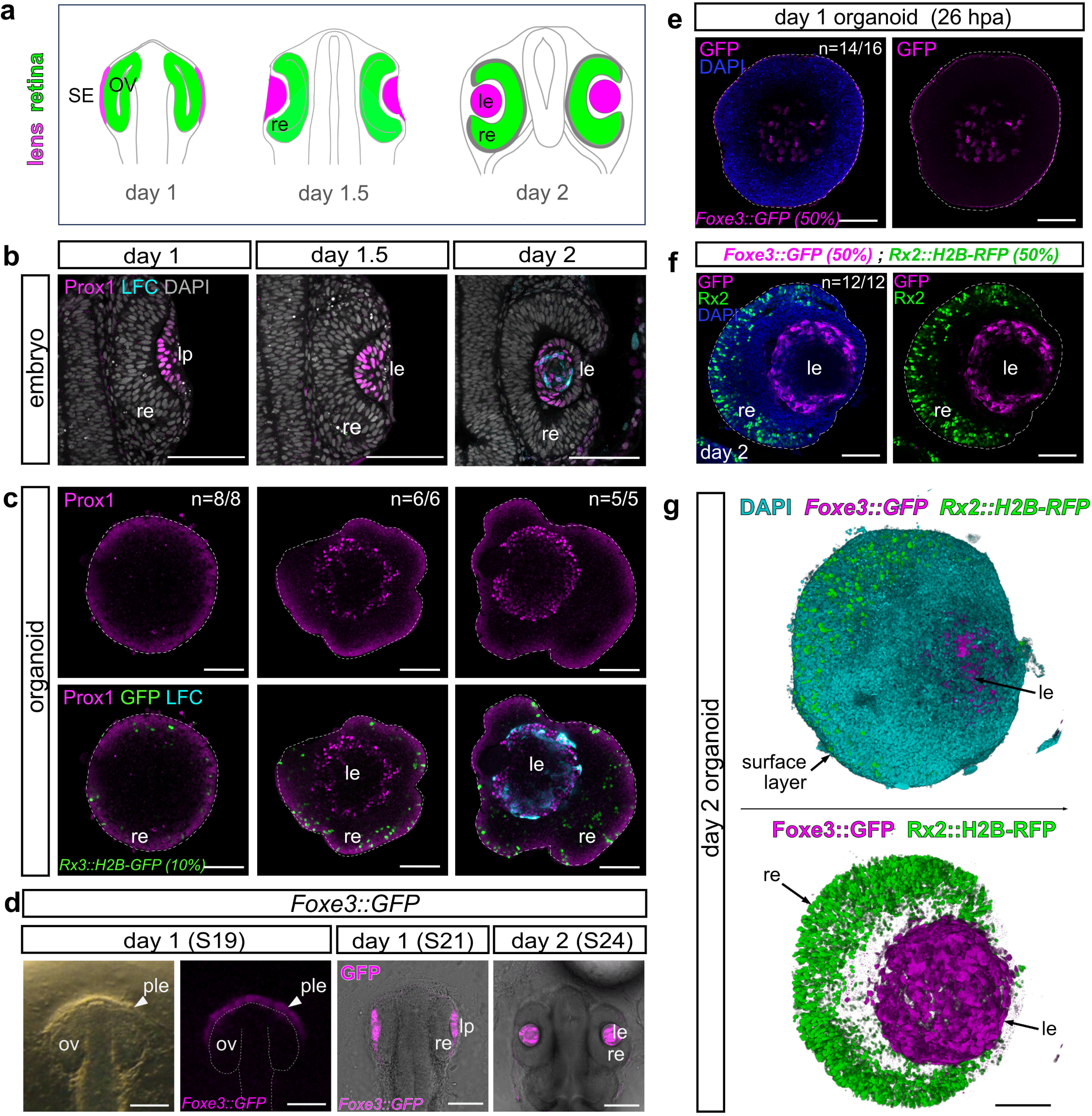
The lens originates from a spatially separated population of lens progenitors in center of the organoid. (See also Figure S2 and S3, Video S1, S3 and S4.) **(a)** Schematic representation of lens formation in medaka. The lens develops from lens-competent head surface ectoderm (SE, magenta), retinal cells from the OV (green). Surface ectoderm forms the lens placode (lp) that further invaginates together with the OV to form the optic cup with a lens sphere surrounded by the retina. **(b)** Optical sections through wild-type medaka embryos at indicated stages stained with the antibody against Prox1, a lens placode-specific marker and LFC, a lens fiber cell marker. **(c)** Optical sections through organoids derived from the *Rx3::H2B-GFP* reporter line stained with antibody against Prox1 (magenta) and LFC (cyan). **(d)** Lens specific GFP expression in embryos of the *Foxe3::GFP* reporter line (magenta). Bright-field and GFP fluorescence images showing reporter expression in presumptive lens ectoderm (ple) of day 1 (S19), lens placode (pl) of 1 dpf (S21) and lens (le) of 2 dpf (S24) medaka embryos. **(e)** Optical section through a day 1 organoid (26 hpa) derived from *Foxe3::GFP* reporter line showing the lens-specific expression of GFP (magenta) detected by anti-GFP antibody, co-stained with DAPI (blue). **(f)** Optical section through a day 2 organoid derived from 50 % *Foxe3::GFP* and 50 % *Rx2::H2B-RFP* reporter blastula cells. Expression of RFP (green) and GFP (magenta), detected by antibodies is mutually exclusive, co-stained with DAPI (blue). **(g)** 3D rendering of *Rx2::H2B-RFP* (green), *Foxe3::GFP* (magenta) organoid shown in f, DAPI stained nuclei are in cyan. n numbers in c and e indicate the number of analyzed organoids with described phenotype. LFC, lens fiber cell; le, lens; re, retina; lp, lens placode; SE, surface ectoderm; OV, optic vesicle; dpf, day post fertilization; hpa, hours post aggregation. Dashed line in c, e and f indicates the outline of the organoid. Dashed line in d indicates the outline of the retina. Scale bar 100 μm.

To specifically address the dynamics, onset, and cellular re-arrangements of lens-committed progenitors, we generated a *Foxe3::GFP* reporter line. *Foxe3* (winged helix/forkhead domain transcription factor) is necessary for proper lens development and is expressed by the cells of the presumptive lens ectoderm and lens epithelium of different species, including medaka (Blixt et al., 2000; Brownell et al., 2000; Mikula Mrstakova and Kozmik, 2024; Shi et al., 2006). This pattern is fully recapitulated by *Foxe3::GFP* with an onset of expression starting at 1 dpf (already in the presumptive lens ectoderm at stage S19 prior to lens placode formation at stage S21 where it is prominently expressed; Iwamatsu et al., 2004). It was further expressed by the majority of lens cells at 2 dpf, (Figure 3d) where it preceded the expression of the lens fiber cell marker LFC (Figure 3-figure supplement 2a). During development, the lens is composed of anteriorly located lens epithelium (proliferative lens progenitors) and posteriorly and centrally located differentiating lens fiber cells (Greiling et al., 2010). We could differentiate those parts of the developing lens with *Foxe3::GFP* expression in both, lens fiber cells and lens epithelium in 3 dpf embryo, and the expression of Prox1 and LFC restricted to differentiating fiber cells, but excluded from the lens epithelium (Figure 3-figure supplement 2a, arrows). Analysis at later stages showed that at 4 dpf, *Foxe3::GFP* expression was maintained in the lens epithelium (proliferatively active progenitors) and was nearly undetectable in differentiated lens fiber cells (Figure 3-figure supplement 2b, arrows).

In organoids, lens progenitor cells expressing *Foxe3::GFP* were first found in the central core at day 1 (26 hours post aggregation) (Video S2, Figure 3e, Figure 4, Video S3). These cells appeared in a pattern complementary to the retinal marker *Rx3::H2B-GFP* (Video S1), indicating that lens progenitor cells originate from a spatially distinct population in the central core of the organoid. We never detected any *Rx3*-positive cells eventually acquiring a lens fate. To address the distribution of lens and retina domains, we generated organoids from mixed blastula cells derived from the retina-specific reporter line *Rx2::H2B-RFP* (Inoue and Wittbrodt, 2011) and the lens specific reporter line *Foxe3::GFP* (Figure 3-figure supplement 2c). By day 2, lens progenitor cells expressing *Foxe3::GFP* contributed to the formation of a spherical lens in the central core of the organoid surrounded by unlabeled cells and eventually by the cortical retinal tissue (Figure 3g, Video S4).

**Figure 4:**
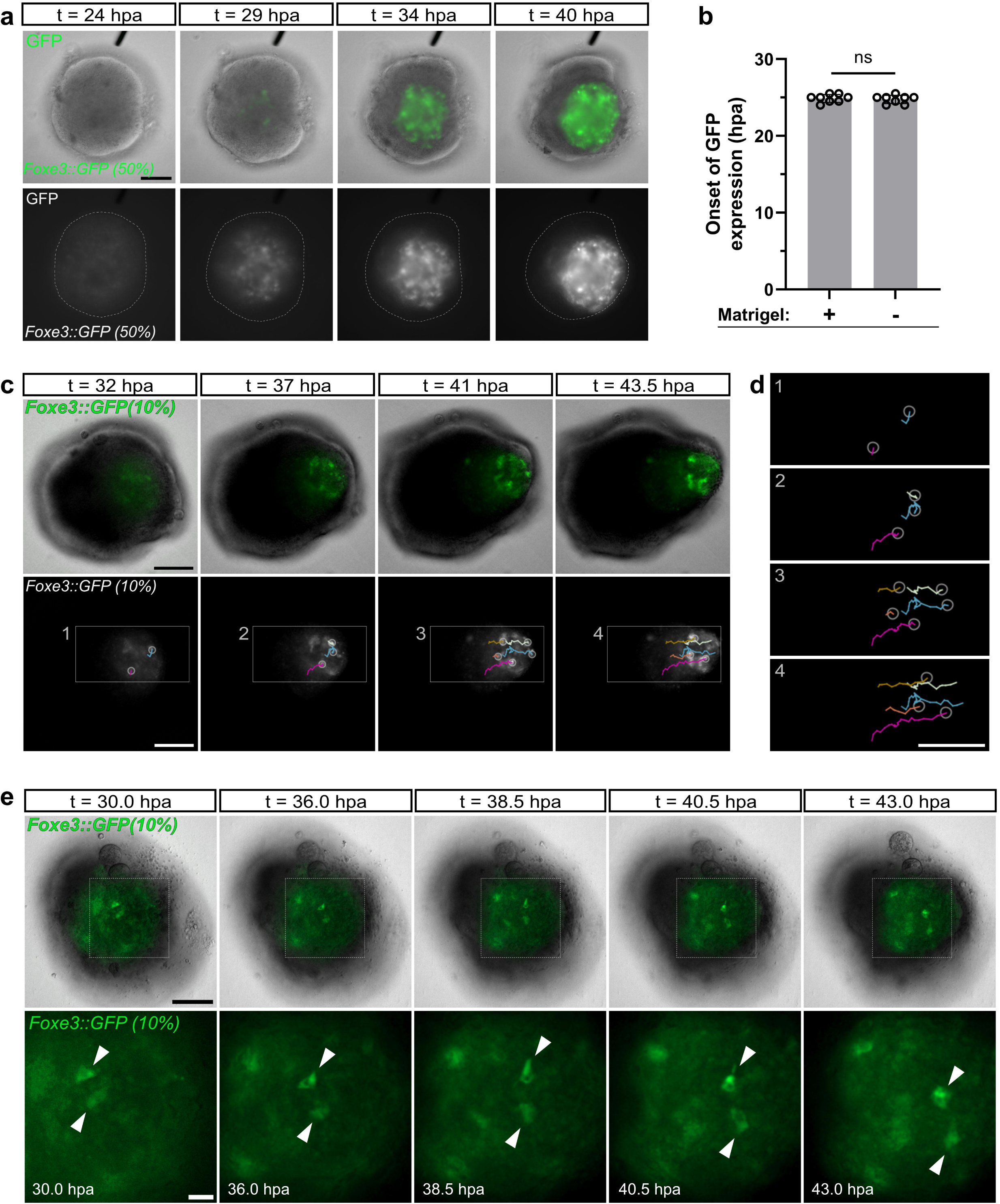
Dynamics of lens formation in ocular organoids. **(a)** Single organoid (from Video S2) generated from *Foxe3::GFP* reporter (50 %) and wild-type blastula cells imaged from day 1 (24 h post aggregation; hpa) to day 2 (40 h post aggregation; hpa). Overlay (top panel) of bright-field and GFP fluorescence and GFP fluorescence (bottom panel, gray) of indicated timepoints between day 1 and day 2 organoid development. Dashed lines indicate the outline of the organoid. Total number of n=62 organoids in 4 independent experiments were followed. **(b)** Onset of GFP expression in *Foxe3::GFP*-derived organoids grown with or without Matrigel supplementation at day 1, n=8 organoids per condition. Statistical significance was calculated using an unpaired two-tailed t-test. n.s., not significant. **(c)** Tracking of individual GFP^+^ cells in the organoids derived from mixture of *Foxe3::GFP* (10 %) and wild-type (90 %) cells from day 1 to day 2. Time indicated in hours post aggregation (hpa). **(d)** Tracks of individual cells over time from details 1-4 in c. **(e)** Movement-associated cell shape changes of GFP^+^ cells in in the organoids derived from mixture of *Foxe3::GFP* (10 %) and wild-type (90 %) cells from day 1 to day 2. Scale bar 100 μm; enlargement in e: scale bar 25 μm.

Further analysis showed that the organoid lens consists of proliferating as well as differentiating cells (Figure 3-figure supplement 2d and e), however not localized to distinct domains as in the embryo. We found that *Foxe3-*driven GFP was co-expressed with the LFC differentiation marker in day 2 organoids (Figure 3-figure supplement 2e). EdU incorporation also showed that the lens is formed by proliferative (EdU^+^) *Foxe3::GFP*-expressing cells (Figure 3-figure supplement 2d). On the surface of day 2 organoid lenses, we initially detected stretches of Foxe3^+^/LFC^-^ cells, reminiscent of the lens progenitor cells of the lens epithelium (Figure 3-figure supplement 2e and f, arrows). However, at day 4, organoid lenses did not show any apparent lens epithelium (Prox1^-^/LFC^-^) (Figure 2g and 2h). In fact, we did not detect any Prox1^-^/LFC^-^ cells on the surface of the day 4 lenses, indicating that in organoid lenses a lens epithelium is absent at later stages.

### Dynamics of lens formation

Analysis of the cellular composition of day 1 and day 2 organoids showed a stereotypic arrangement of cell types: retinal-committed progenitors established in the epithelialized surface while cells localized in the core of the organoid adopted lens fate and formed centrally localized lens spheres (Figure 3-figure supplement 1d, Figure 3c and 3e). Many morphogenetic events such as germ layer segregation, boundary sharpening, and compartmentalization can be at least partly explained by cell sorting driven by adhesive and mechanical differences (Amack and Manning, 2012; Krieg et al., 2008; Steinberg, 1963; Zhang et al., 2011). To address whether differential adhesive properties of lens and retinal progenitor cells contribute to such cellular arrangements, particularly to the arrangement of lens-committed progenitors into a lens sphere, we performed dissociation/re-aggregation experiment (Figure 4-figure supplement 1a). Day 1 organoids originating from the *Foxe3::GFP* reporter line were dissociated into a single cell suspension to disrupt presumptive retinal/lens domains and to ensure a random distribution of lens-/retina-committed cells (Figure 4-figure supplement 1b-c). Those suspensions were subsequently allowed to re-aggregate (Figure 4-figure supplement 1d, Video S5). Time-laps imaging showed a gradual sorting of *Foxe3::GFP* labeled cells – from single GFP^+^ cells (Figure 4-figure supplement 1c) through groups of GFP^+^ cells (after 3h, Figure 4-figure supplement 1d, Video S5) to one single lens localized in the center of the organoid by day 2 (15 h), surrounded by retinal neuroepithelium (AcTub^+^ cells) (Figure 4-figure supplement 1d and e), scenario resembling organoid organization before dissociation. This data show that differential adhesive properties of lens/retinal precursor cells can enable formation of a compact spherical lens in the center of the organoid.

Although lens progenitors were initially established in the center of the day 1 organoid, the analysis of day 2 and day 4 organoids indicated that the lens was displaced from this initial position to reach the periphery of the organoid by day 2 (Figure 3g, Video S4 and Figure 2b). This was confirmed by time-lapse imaging of ocular organoid development from day 1 to day 2 where *Foxe3::GFP*-expressing cells appear initially in the core region of the organoid at 25 hours post aggregation where they organize into a spherical lens within 16 hours (Video S2, Figure 4a and 4b). This process was accompanied by a gradual shift of the forming lens from center to the periphery where the lens is eventually reaching the surface of the organoid (Video S2, Figure 3f and 3g, Figure 4a). Tracking of individual lens-committed cells supported this trend. Individual centrally located cells collectively moved to the organoid periphery (Figure 4c and 4e, Video S6). This process was accompanied by profound changes in cell shape of tracked *Foxe3::GFP* cells (Figure 4d, Video S7) indicating an active movement of lens progenitors to the organoid periphery. On the other hand, nuclei of retinal cells (*Rx3::H2B-GFP*) followed over same period of time, demonstrated short oscillating movements confined to the retinal neuroepithelium on the surface of the organoid (Video S8), representing interkinetic nuclear migration found in differentiating retinal progenitor cells within the retina. Taken together, these data indicate that differential adhesion-based cell sorting between lens and retinal progenitors together with an active movement of lens progenitor cells from the center to the periphery of the organoid are the driving force of the lens sphere formation and its displacement through the static retinal epithelium.

### Lens size scales with organoid size

Since retinal and lens progenitors occupied distinct regions within the organoid, we next investigated how their distribution is affected by total organoid size. We generated organoids from different numbers of cells at day 0 (4,000, 2,000, 1,000 and 500 cells). This approach resulted in organoids of different sizes at day 1: organoids derived from 4,000 cells had an average diameter of 521 ± 13 µm, those from 2,000 cells measured 379 ± 15 µm, from 1,000 cells 310 ± 22 µm, and from 500 cells 201 ± 55 µm (Figure 5a, 5b and 5d). Interestingly, the thickness of the tissue acquiring retinal progenitor cell identity in the cortex of the ocular organoid (*Rx3::H2B-GFP*) remained constant, independent of the initial size of the aggregate reaching to the depth of: 59.7 μm ± 1.6 μm for 4,000 cells; 55.6 ± 3.1 μm for 2,000 cells; 56.4 μm ± 3.2 μm for 1,000 cells and 55.4 μm ± 6.8 μm for 500 cells (Figure 5b and 5e). While the thickness of retinal neuroepithelium stayed constant, the lens size scaled proportionally with the size of the organoid. By day 4, larger organoids developed larger lenses, whereas smaller organoids formed smaller lenses (224 μm ± 12 μm for 4,000 cells; 199 μm ± 26 μm for 2,000 cells; 115 μm ± 33 μm for 1,000 cells and 68 μm ± 20 μm for 500 cells) (Figure 5c and 5f). Although 500 cell derived organoids were able to form lenses, the efficiency of lens formation was considerably lower (33.3 %; Figure 5g) than in organoids of the bigger sizes (between 86.7 - 95 % for organoids generated by 1,000 or more cells) (Figure 5g). Ocular organoids generated by the aggregation of less than 500 cells did not form any lenses (Figure 5g). Taken together, these data indicate that lens size scales with the size of the organoid and is defined by the constant thickness of the forming retina.

**Figure 5.**
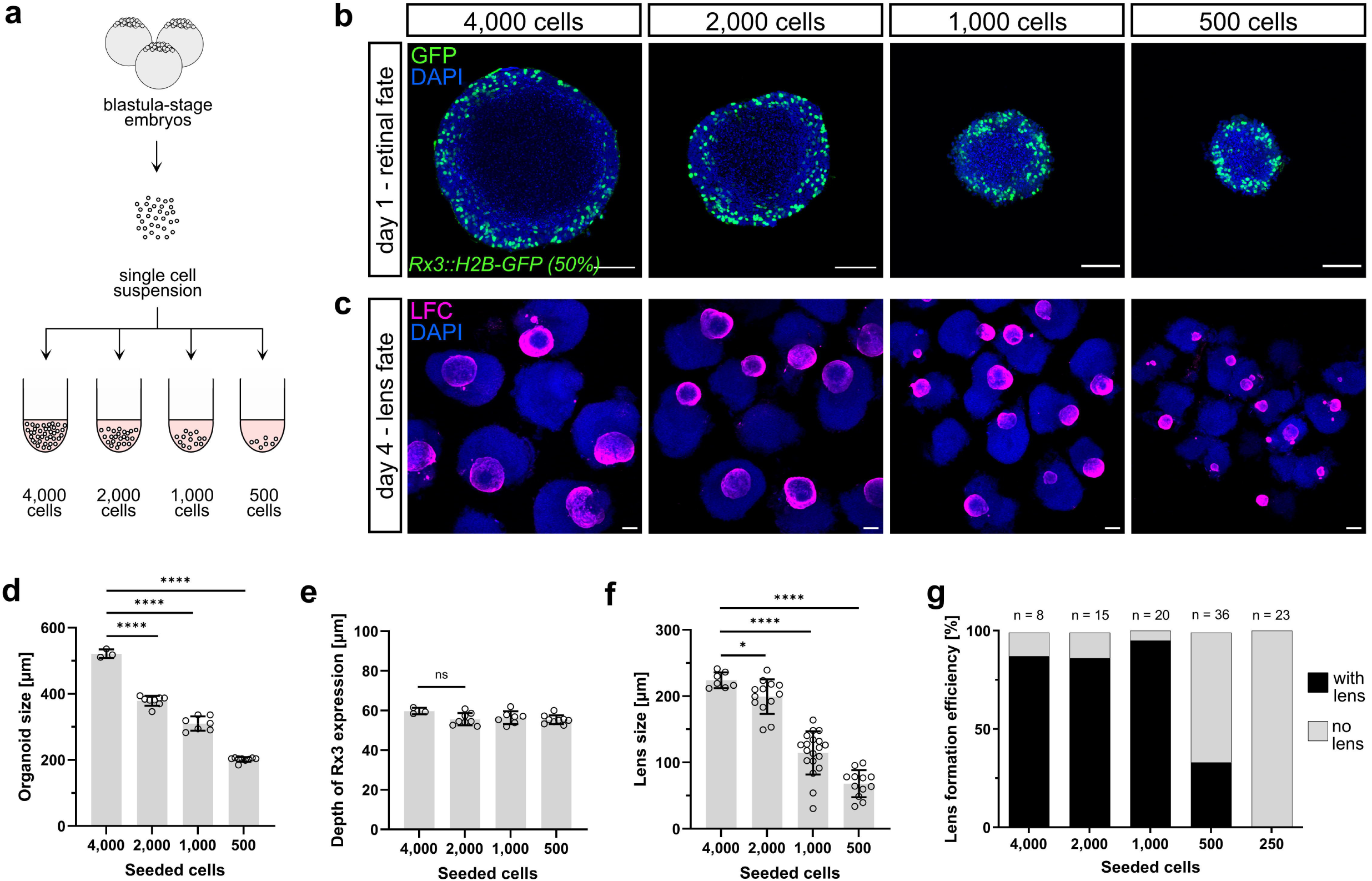
Lens size scales with organoid size. **(a)** Schematic representation of generation of organoids of different sizes. Pluripotent embryonic cells were seeded in different seeding densities. **(b)** Representative optical sections of day 1 organoids generated by aggregation of indicated number of cells from *Rx3::H2B-GFP* reporter line immunolabeled with anti-GFP antibody, co-stained with DAPI. For *Rx3::H2B-GFP*, the percentage (%) indicates how many cells carry the indicated transgene. **(c)** Representative images of day 4 organoids generated by indicated number of cells stained with an anti-LFC antibody (maximal z-projections). **(d)** Quantification of diameter of day 1 organoids generated by aggregation of indicated number of cells. **(e)** Quantification of depth of retina-specific marker expression (*Rx3::H2B-GFP*) at day 1 generated by aggregation of indicated number of cells. Number of analyzed organoids: n=3 for 4,000 cells, n=8 for 2,000 cells, n=7 for 1,000 cells, n=9 for 500 cells. **(f)** Quantification of lens size measured as diameter of the lens in day 4 organoids. n=7 for 4,000 cells, n=13 for 2,000 cells, n=19 for 1,000 cells, n=19 for 500 cells. **(g)** Efficiency of lens formation in organoids of different sizes measured as proportion of organoids carrying lens judged by immunolabeling with anti-LFC antibody. Statistical significance was calculated using an unpaired two-tailed t-test. n.s., not significant; ****p<0.0001, *p<0.05. le, lens; re, retina; LFC, lens fiber cell. Scale bar 100 μm.

### Lens formation depends on temporal activity of FGF and BMP signaling

The competence to form a lens is established during the formation of the anterior neural plate within the region of the anterior neural plate border and is regulated by the coordinated activity of BMP and FGF signaling (Gunhaga, 2011; Litsiou et al., 2005; Sjödal et al., 2007). Once lens competence is established within the SE, FGF and BMP signaling induce its differentiation into the lens placode and subsequently into the lens, accompanied by the expression of structural lens proteins (Faber et al., 2001; Furuta and Hogan, 1998; Le and Musil, 2001; Rajagopal et al., 2009; Sjödal et al., 2007; Wawersik et al., 1999; Zhao et al., 2008). Both BMP and FGF signaling pathways were found to be activated in lens-forming organoids. We used specific antibodies to detect the phosphorylated form of SMAD1/5/8 (pSMAD1/5/8) and ERK kinase (pERK1/2) as downstream effectors of BMP and FGF signaling, respectively (Figure 6a and 6b). BMP signaling activity was detected in the center (region of establishment of lens-committed progenitors (Figure 3e)) of the organoid at day 1 (Figure 6a). On the other hand, only sparse FGF signaling activity (as visualized by pERK1/2) was detected in day 1 organoids, with activity peaking later - at day 1.5 in the center of the organoid. At day 2, FGF activity was observed in both, retinal and lens cells corresponding to the FGF activity in the developing embryo (retina and lens epithelium) at corresponding stage (Figure 6b). To address the contribution of the observed BMP and FGF signaling activity to organoid lens formation, we exposed developing organoids to FGF and BMP antagonists, SU5402 and Noggin or Dorsomorphin, respectively. To separate the establishment of lens competence from the process of lens differentiation, we divided the treatments into two discrete time windows: day 0 to day 1 corresponding to the time before the lens placode formation (establishment of lens competence), and day 1 to day 2 covering lens placode formation and differentiation of lens fiber cells (Figure 6c). We then assessed the organoids at day 2 for the expression of the early lens progenitor marker *Foxe3::GFP* and the differentiation marker LFC.

**Figure 6.**
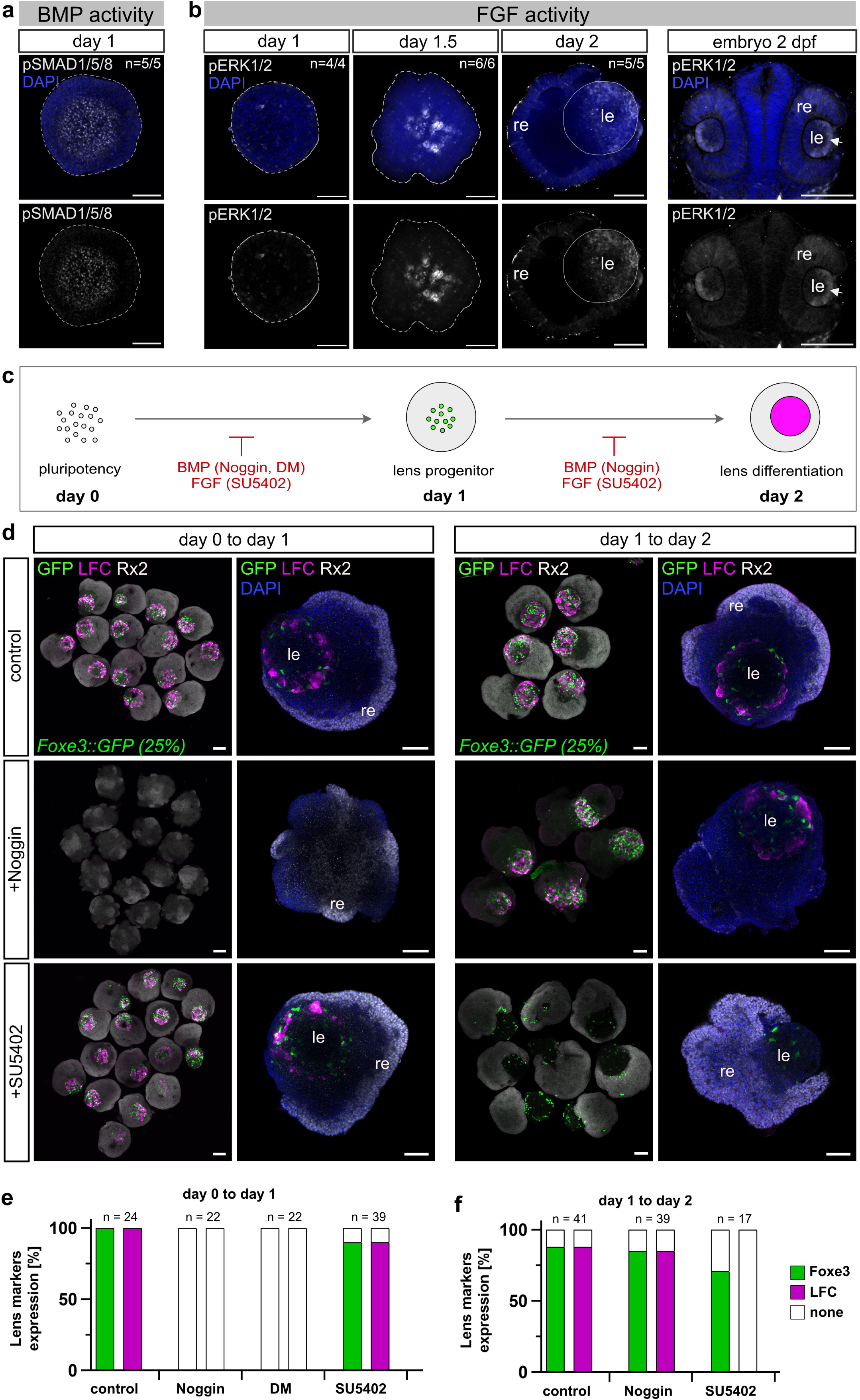
Lens formation is dependent on temporal activity of FGF and BMP signaling. **(a)** Activity of BMP signaling detected by immunohistochemistry against phosphorylated SMAD1/5/8 (pSMAD1/5/8) in day 1 organoids. **(b)** Activity of FGF signaling detected by immunohistochemistry against phosphorylated ERK1/2 (pERK1/2) at indicated stages of organoid development and in embryo at 2 dpf. Dashed line indicates the outline of the organoid in day 1 and day 1,5 organoid. In day 2 organoid, line outlines the lens. n numbers indicate the number of organoids with displayed phenotype within one experiment. **(c)** Schematic representation of organoid treatment with FGF and BMP signaling antagonists. Organoids were generated from *Foxe3::GFP* reporter line. Inhibitors of BMP (Noggin, 100 ng/μl or Dorsomorphin -DM, 100 ng/ml) or FGF (SU5402, 10 μM) signaling were administered in time window from day 0 to day 1 or from day 1 to day 2. Organoids were analyzed at day 2 for the expression of lens progenitor marker *Foxe3::GFP* and lens fiber cell differentiation marker LFC. **(d)** Representative images of day 2 organoids treated as indicated and labeled with antibodies against GFP (green), LFC (magenta) and Rx2 (grey). Overview images are represented by maximal z-projections; single organoids are represented by single optical sections. For *Foxe3::GFP*, the percentage (%) indicates how many cells carry the indicated transgene. **(e-f)** Quantification of expression of *Foxe3::GFP* and LFC in control and treated organoids measured as proportion of organoids expressing lens progenitor marker *Foxe3::GFP* (green) and lens differentiation marker LCF (magenta). Total number of n=204 organoids grouped in control and treated groups were analyzed. All treatments were performed in two biological replicates. re, retina; le, lens; DM, Dorsomorphin; LFC, lens fiber cell. Scale bar 100 μm.

Blocking BMP signaling with Noggin or Dorsomorphin completely prevented the expression of lens-specific markers if applied from day 0 to day 1 (Figure 6d and 6e) but did not impact on lens formation if administered from day 1 to day 2 (Figure 6d and 6f). In contrast, inhibition of FGF signaling with SU5402-mediated did not affect initial lens formation if applied from day 0 to day 1 and allowed expression of both, lens placode and lens differentiation markers. However, it efficiently blocked the differentiation of lens fiber cells when applied from day 1 to day 2, and completely prevented the expression of the LFC marker (Figure 6d, 6e and 6f). This indicates that similar to embryonic development, the establishment of lens competence (in the center of the organoid between day 0 and day 1) depends on the activity of BMP signaling, while FGF signaling is required later for the lens differentiation process (day 1 to day 2).

### Lens organoid gene expression recapitulates the onset of lens-specific genes *in vivo*

To address the temporal dynamics of lens-specific gene expression in *in vivo* and in organoid-based lens development, we performed total RNA sequencing analysis. Embryos at 1, 2, 3, and 4 dpf as well as organoids at corresponding developmental stages, were analyzed for the expression of key markers of lens development. For total RNA sequencing we extracted RNA from whole organoids (composed of both lens and retina), the anterior part of day 1 and 2 embryos (dissected right behind optic vesicles or optic cups), and the eyes of 3 dpf and 4 dpf embryos (Figure 7a). Within the total RNA reads we focused specifically on key genes implicated in the different stages of lens development. Among the genes analysed, the earliest expressed was the transcription factor *Foxe3*, with expression peaking at day 1 of development, followed by the upregulation of the transcription factor *Pitx3* (Figure 7c, Data S1). Day 2 marks the onset of expression of main structural lens proteins in embryos and organoids, namely: crystallins: *cryaa* (αA-crystallin), *cryba1a* (βA1-crystallin), *crybb1l2* (βB1 like 2), *crybb1l3* (βB1 like 3), *cryba2b* (βA2), *cryba4* (βA4), *crybb1* (βB1), *crygn2* (γN2-crystallins), *crygmx* (γMX), *crygm3* (γM3); major lens membrane structural proteins (MIPs/acquaporins) *mipa* and *mipb* coding for water channels ensuring water transport between lens cells (Agre and Kozono, 2003; Donaldson et al., 2024; Varadaraj et al., 1999); Gap junction proteins *Gja8* (connexin-50) and *Gje1* (connexin-23) and lens intrinsic membrane proteins *lim2.2* and *lim2.4* facilitating intercellular transport of metabolites and ions in lens fiber cells (Mathias et al., 2010). Day 3 was characterized by the onset of expression *hsf4* (heat shock transcription factor 4), coding for a transcription factor required for de-nucleation and organelle degradation during lens terminal differentiation (Cui et al., 2013; Gao et al., 2017). Accordingly, *dnase1l1l* (deoxyribonuclease I-like 1-like) and *plaat1* (phospholipase A/acyltransferase) implicated in DNA and lens organelle degradation, respectively (Morishita et al., 2021; Zhang et al., 2020) were also expressed from day 3 of development. At the same stage, the expression of proteins making up the structural scaffold critical for fiber cell architecture and mechanical properties of the lens was initiated. Those include beaded filament proteins *Bfsp1* (filensin) and *Bfsp2* (phakinin) and *Lgsn* (lengsin) (Harding et al., 2008; Perng et al., 2007; Song et al., 2009).

**Figure 7.**
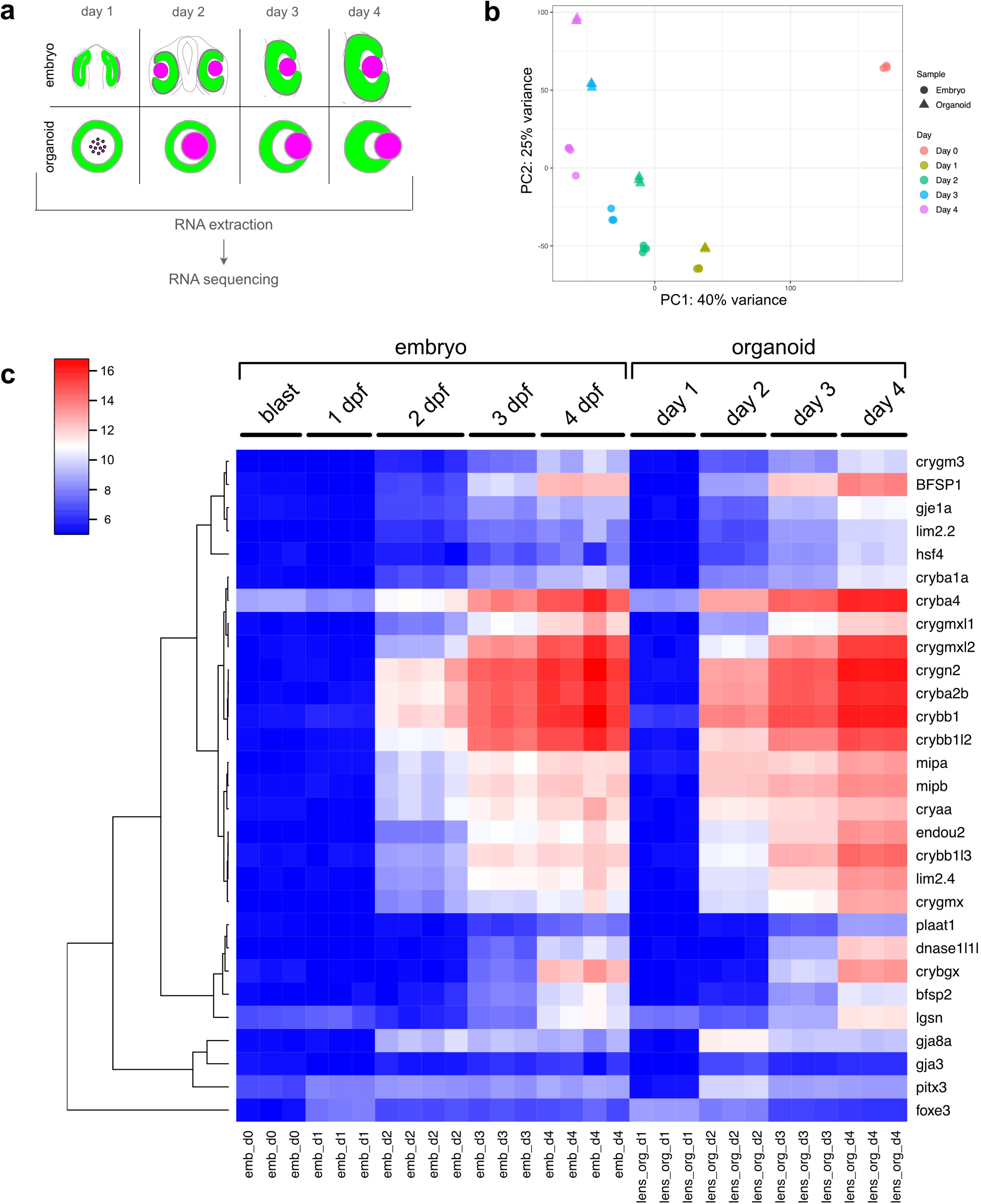
Lens organoid gene expression recapitulates gene expression *in vivo*. **(a)** Schematic representation of sample preparation. Whole organoids (composed of both lens and retinal tissue) were analyzed. For 3 dpf and 4 dpf medaka embryos, eyes were dissected (composed of both lens and retina). For 1 dpf and 2 dpf embryos the anterior part of the head right behind optic vesicles or optic cup, respectively was dissected and used for RNA isolation. **(b)** Principal component analysis of the transcriptomes of embryonic and organoid samples at the indicated stages. **(c)** Heatmap clustering of variance stabilized counts of lens-specific genes. Samples were processed in 3 technical replicates each. blast, blastula-stage embryo; emb, embryo; lens_org, lens organoid.

Our comparative analysis indicates highly reminiscent gene expression patterns of key marker genes when comparing organoid and embryonic lens development (Figure 7c, Data S1). This demonstrates that our conditions promote the development of ocular organoids in which the expression of key markers genes is temporally recapitulating the order of expression in the development of the embryonic lens.

## Discussion

*In vitro* organoid development builds on the remarkable ability of embryonic stem cells or progenitor cells to autonomously self-organize in higher order structures with sophisticated microarchitecture, often closely resembling cell type compositions, 3D structure, and the function of the organ (Camp et al., 2015; Dye et al., 2015; Eiraku et al., 2011; Lancaster et al., 2013; McCracken et al., 2014; Nakano et al., 2012).

Here, we used medaka derived pluripotent cells as a model for the generation of complex ocular organoids composed of both, retina and lens. We showed that fish retinal organoids efficiently form embedded lenses, in a process that employs the molecular pathways including expression of key transcription factors and utilization of extracellular signaling pathways, as the embryo. In contrast to *in vivo* development, organoid cells did not follow the canonical process of lens formation, but used an alternative mode of cellular assembly to generate and grow the lens. While in the embryo, the lens sphere is shaped by the coordinated invagination of lens committed head surface ectoderm and retina-forming optic vesicle neuroepithelium, lens progenitors in the organoid were established in a different fashion directly inside of the retinal organoid to form the lens.

By following early progenitors of both retina (*Rx3*) and lens (*Foxe3*) we found that retina progenitor identity was acquired at the surface, while a spherical domain with lens progenitor identity was established in the center of the organoid (Figure 3c and Figure 3e). While in the embryo the lens originates from the head surface ectoderm, non-canonical routes to lens regeneration have been described in some species. In newts, the lens can be regenerated from the pigmented epithelial cells of the iris after injury or lens removal (Agata et al., 1993; Grogg et al., 2005; Maki et al., 2007). Lens formation was also observed in the cultures of chick neuroretinal cells (Iida et al., 2017; Okada et al., 1975) showing that retinal-fated cells can give rise to lens cells under certain circumstances. However, in medaka ocular organoids, retina and lens are formed by spatially separated populations of retina- and lens-committed progenitor cells.

The size of the lens scaled with the size of the organoid and was limited by the volume of the organoid that acquired retinal fate (Figure 5). The constant depth of retinal fated cells in the cortex under the surface of the organoid, regardless of the size of the aggregate, suggests that retinal fate is influenced by environmental gradients, which cause the diffusion-limited availability of retina-inducing factors. It has previously been shown that cells on the surface of the organoid experience higher level of oxygen tension, higher level of nutrients and signaling molecules than cells located more centrally (Grün et al., 2023; McMurtrey, 2016; Qian et al., 2019; Tse et al., 2021) when grown in 3D suspension culture. Additionally, cells at the surface experience different mechanical forces, mostly due to the interactions with ECM (Vining and Mooney, 2017). Both, retinal as well as lens fates, were established either before ECM addition or in the absence of Matrigel (Zilova et al., 2021, this study), making the environmental gradient more likely to contribute to the observed tissue arrangement in fish organoids.

Interestingly, Bmp inhibition in the early stages of organoid development, before the appearance of *Foxe3*-expressing lens progenitor cells (day 0 to day 1) completely abolished lens formation, while BMP inhibition during the lens formation process (from day 1 to day 2) did not show any impact. Thus, we hypothesize that a source of BMP in the forming neuroretina establishes initial lens competence in the center of the organoid. During normal development, BMP4 is secreted by the neuroepithelium of the developing OV and was found to be essential for lens induction (Furuta and Hogan, 1998; Rajagopal et al., 2009). Accordingly, ablation of the prospective neuroectoderm in chick (Kamachi et al., 1998) or defective OV formation in mouse (Klimova and Kozmik, 2014; Mathers et al., 1997; Porter et al., 1997) prevents lens formation, indicating that the developing neuroepithelium stimulates lens formation within the pre-placodal ectoderm. The emergence of lens fate in the center of ocular organoids consequently depends on a critical threshold of the morphogen to be reached in the center. The scaling of lens size with organoid size supports this hypothesis as larger organoids are better suited to reach this threshold - potentially due to increased total ligand production (in the forming neuroretina) resulting in the formation of a more pronounced gradient.

During normal development, developing retinal cells have been suggested to serve as a source of FGFs stimulating expression of lens structural genes and inducing lens differentiation (Chow et al., 1995; Faber et al., 2001; Gunhaga, 2011; Le and Musil, 2001; McAvoy et al., 1999; Robinson, 2006; Zhao et al., 2008). Accordingly, blocking of FGF signaling activity during organoid lens formation prevented lens differentiation. The dependency of lens formation on the activity of BMP and FGF signaling pathways has been well documented also in other *in vitro* systems where transient activation of BMP and FGF signaling is necessary for the *in vitro* differentiation of lens-like structures (lentoid bodies) (Dincer et al., 2013; Fu et al., 2017; Leung et al., 2013; Yang et al., 2010).

Since organoids show remarkable similarities to their *in vivo* counterparts when it comes to cellular composition and key structural features, we naturally tend to infer that corresponding structures are formed by corresponding cellular processes in both systems. Here we show an example of an alternative mode of cellular assembly that leads to a similar structural and molecular outcome. In the embryo, the lens formation is driven by invagination of the lens placode to generate a lens pit and finally a spherical lens. In organoids on the other hand, individual lens-committed progenitors are established in the center of the organoid. Our current data support the hypothesis, that differential adhesive properties of lens and retinal progenitor cells mediated by preferential homotypic cell-cell adhesion support the formation of the lens sphere. Concomitant active movement of lens-committed progenitors from the organoid center to its periphery through the retinal sheet on the surface places the lens to the superficial position, resulting in the observed optic cup-like morphology of the organoid. Our observation that dissociated organoid cells re-aggregate into homotypic clusters is consistent with classical and more recent work showing that differential adhesion, often combined with differential contractility, can drive robust cell sorting in both embryonic tissues and *in vitro* systems. Such adhesion-based sorting is now regarded as a generic mechanism that frequently contributes to tissue assembly and compartmentalization during embryonic development, alongside complementary processes such as active cell migration and contact inhibition of locomotion (Amack and Manning, 2012; Brayford et al., 2019; Davis et al., 2012; Krieg et al., 2008; Zhang et al., 2011).

Organoids can recapitulate many aspects of embryonic development. However, the absence of embryonic constraints can also lead to significant differences in tissue organization, cellular interactions, and overall structure. While the goal of many organoid studies is to replicate *in vivo* conditions as closely as possible (Chen et al., 2024; Hofer and Lutolf, 2021; Kretzschmar and Clevers, 2016; Sahu and Sharan, 2020; Zhao et al., 2022), the unconstrained environment of organoids offers a unique opportunity to tap the organoid’s potential of self-organization. This can uncover alternative developmental pathways that are not apparent *in vivo* and could expand our understanding of the plasticity, robustness and adaptability of developmental programs when unleashed from evolutionary constraints. Insights gained from alternative assembly processes could inform the design of synthetic biological systems and potentially lead to innovative strategies for creating complex tissues and organs *in vitro*.

## Materials and Methods

### Fish handling and maintenance

Medaka (*O. latipes*) stocks were maintained according to the local animal welfare standards (Tierschutzgesetz §11, Abs. 1, Nr. 1, husbandry permit AZ35-9185.64/BH, line generation permit number 35–9185.81/G-145/15 Wittbrodt). The following medaka lines were used in this study: Cab strain as a wild type (Loosli et al., 2000), *Rx3::H2B-GFP* (Rembold et al., 2006), Rx2::H2B-GFP (Inoue and Wittbrodt, 2011), *Atoh7::EGFP* (Del Bene et al., 2007), *Gaudí^RSG^* (Centanin et al., 2014), *Foxe3::GFP* (this study).

### Generation of organoids

Organoids were generated as previously described (Zilova et al., 2021) with slight modification of the protocol. Blastula stage embryos, stage 11 (Iwamatsu, 2004) were de-chorioneated, cells mass was isolated and washed twice with PBS (Thermo Fisher Scientific, Cat#:10010023). Cell mass was dissociated in PBS by pipetting up and down and centrifuged through 40 μm strainer (pluriSelect, Cat#:43-10040-51) for 3 min at 180 x g. Pellet was re-suspended in differentiation media: DMEM/F12 (Gibco, Cat#:11320033), 5 % KSR (Gibco, Cat#:10828028), 1x NEAA (Sigma-Aldrich, Cat#:M7145), 1x sodium pyruvate (Sigma-Aldrich, Cat#:S8636), 1x penicillin/streptomycin, 20 mM Hepes pH 7.4 (Roth, Cat#:9105.4), 0.1 μM ϕ3-mercaptoethanol (Gibco, Cat#:21985-023). If not stated otherwise, 3.000 cells per aggregate in 100 μl were seeded into individual wells of low-binding 96-well plate (Thermo, Cat#:174925 Nunclon Sphera). Cells were incubated in 26°C in the incubator without CO_2_ control. At day 1, aggregates were washed in differentiation media, Matrigel (Corning, Cat#:356238) was added to final concentration of 2 % and aggregates were incubated in humidified tissue culture incubator under 5 % CO_2_ condition.

To label specific population of cells, organoids were generated from fish reporter lines expressing fluorescent protein under the control of tissue specific promotor/enhancer (*Cry::ECFP*; *Rx3::H2B-GFP*; *Rx2::H2B-RFP*; *Atoh7::EGFP*; *FoxE3::GFP*). In that case, cells were extracted from the blastula stage embryos originating from the outcross of fish reporter line and wild-type fish, and in some cases further mixed with cells originating from wild-type embryos of the same stage to achieve sparse labeling. The proportion of cells coming for transgenic reporter line genetically able to express the transgene is indicated in the figures in percentage (%).

### Treatment of cells/aggregates/organoids

Cells or aggregates were treated with Noggin (Merck Millipore, Cat#:GF173, 100 ng/ml), Dorsomorphin (Merck, Cat#:171260, 100 ng/ml), or SU5402 (Sigma, Cat#:SML0443, 10 μM). For treatment from day 0 to day 1, blastula stage cells were resuspended directly in differentiation media containing an antagonist in desired concentration. At day 1 (16 hours post aggregation), aggregates were washed twice with fresh differentiation media, supplemented with 2% Matrigel (Corning, Cat#:356238) in differentiation media and incubated till day 2. For treatment from day 1 to day 2, day 1 aggregates (16 hours post aggregation) were washed with differentiation media and transferred into antagonist-containing media and immediately supplemented with 2% Matrigel diluted in antagonist-containing media and incubated till day 2. All organoids were fixed at day 2 and analyzed for the presence of lens and retinal tissues by immunohistochemistry.

### Dissociation and re-aggregation of organoids

Day 1 organoids (25 hours post aggregation) derived from *Foxe3::GFP* reporter line (100% cells labeled) were washed in PBS (Thermo Fisher Scientific, Cat#:10010023) and dissociated in mixture of Trypsin (2.5 %; Gibco, Cat#:15090-046) and Dispase (1 U/ml; STEMCELL Technologies, Cat#:07923) in ratio 1:1. 5-10 organoids we dissociated in 150 μl of dissociation mix for 10 min at 32°C. Dissociation was stopped by addition of 150 μl of 50 % FBS (Gibco, Cat#:10063732) in PBS. Cell suspension was centrifuged through 40 μm strainer (pluriSelect, Cat#:43-10040-51) for 3 min at 180 x g and pellet was re-suspended in differentiation media: DMEM/F12 (Gibco, Cat#:11320033), 5 % KSR (Gibco, Cat#:10828028), 1x NEAA (Sigma-Aldrich, Cat#:M7145), 1x sodium pyruvate (Sigma-Aldrich, Cat#:S8636), 1x penicillin/streptomycin, 20 mM Hepes pH 7.4 (Roth, Cat#:9105.4), 0.1 μM ϕ3-mercaptoethanol (Gibco, Cat#:21985-023) and seeded into wells of low-binding 96-well plate (Thermo, Cat#:174925 Nunclon Sphera). Cells were incubated in 26°C in humidified tissue culture incubator under 5 % CO_2_ condition or imaged with ACQUIFER Imaging Machine (ACQUIFER Imaging GmbH, Heidelberg, Germany) (Pandey et al., 2019).

### Fluorescent labeling

Fish embryos and organoids were fixed in 4% PFA overnight at 4°C, washed with PTW (PBS supplemented with 0.1 % Tween 20), permeabilized by incubation in acetone for 15 min in - 20°C and washed tree times with PTW. Samples were blocked in PTW supplemented with 4 % sheep sera (Sigma, Cat#:14C509), 1 % BSA (Sigma, Cat#:4503) and 1 % DMSO (Sigma, Cat#:D4540) and incubated with primary antibody diluted in 0.1 x blocking solution overnight at 4°C. Primary antibodies used in this study: rabbit anti-Rx2 (Reinhardt et al., 2015) (1:500), chicken anti-GFP (Invitrogen, Cat#:A10262, 1:500), mouse anti-lens fiber cell (Abcam, Cat#:ab185979, ZL-1, 1:500), rabbit anti-Prox1 (Millipore, Cat#:AB5475, 1:2000), rabbit anti-RFP (Clontech, Cat#:632496, 1:500), rabbit anti-pSMAD1/5/8 (Cell Signaling, Cat#: 9511S, 1:500); rabbit anti-pERK1/2 (Cell Signaling, Cat#:4370, 1:500); mouse anti-acetylated tubulin (Sigma, Cat#: T7451, 1:500). Samples were washed with PTW and incubated with the mixture of DAPI nuclear stain (Sigma, Cat#:D9564, 1:500) and secondary antibodies diluted in 0.1 x blocking solution overnight at 4°C. Secondary antibodies used in this study: donkey anti-mouse 647 (Jackson, Cat#:105869, 1:750), donkey anti-chicken 488 (Jackson, Cat#:703-545-155, 1: 750), donkey anti-rabbit 594 (Invitrogen, Cat#:A21207, 1:750). Samples were washed with PTW. For detection of proliferating cells, Click-iT EdU Alexa Fluor 647 Imaging Kit (Invitrogen, Cat#:C10340) was used according to manufacturer instructions. All immunolabeled samples were transferred into clearing solution composed of: 20% urea (Vetec, Cat#:V900119), 30% D-sorbitol (Vetec, Cat#:V900390), 5% glycerol (Vetec, Cat#:V900122) in DMSO (Vetec, Cat#:V900090) (Zhu et al., 2019) and imaged with laser scanning confocal microscope Sp8 (Leica).

### Imaging

Fixed immunolabeled samples were imaged in MatTek glass bottom dish (Mattek, Cat#:D35G-1.5-10-C) with Sp8 confocal microscope (Leica). Organoids overview bright-field pictures were taken with wide-field microscope (Mi8; Leica). For live imaging of lens formation (Video S2, Figure 4a), organoids generated from *Foxe3::GFP* reporter line (50 % of reporter cells mixed with 50 % of wild-type cells) were imaged from day 1 (24 h post aggregation) to day 2 (40 h post aggregation). Imaging was caried out at 26°C directly in low binding 96-well plate (Thermo, Cat#:174925 Nunclon Sphera) with ACQUIFER Imaging Machine (ACQUIFER Imaging GmbH, Heidelberg, Germany) (Pandey et al., 2019). For each well, a set of 10 z-slices (70 µm step size) was acquired in the bright field (50% LED intensity, 40 ms exposure time) and 470 nm fluorescence (50% LED excitation source, FITC channel, 200 ms exposure time) channels. Imaging was performed over 16 hr of organoid development with 30 min intervals. For live imaging of re-aggregation of dissociated cells (Figure 4-figure supplement 1d and Video S5), organoids generated from *Foxe3::GFP* reporter line (100 % of reporter cells) were imaged from day 1 (25 h post aggregation/1 h after dissociation) to day 2 (15 h after dissociation). Imaging was caried out at 26°C directly in low binding 96-well plate (Thermo, Cat#:174925 Nunclon Sphera) with ACQUIFER Imaging Machine (ACQUIFER Imaging GmbH, Heidelberg, Germany) (Pandey et al., 2019). For each well, a set of 30 z-slices (10 µm step size) was acquired in the bright field (50 % LED intensity, 40 ms exposure time) and 470 nm fluorescence (100 % LED excitation source, FITC channel, 50 ms exposure time) channels. Imaging was performed over 15 hr of with 1 h intervals.

For tracking of individual cells (Figure 4c and 4e; Video S6, S7 and S8), organoids generated from *Foxe3::GFP* (10 % of cells from transgene mixed with 90% of cells from wild-type embryos to ensure sparse labeling) or *Rx3::H2B-GFP* (2 % of cells from transgene mixed with 98% of cells from wild-type embryos to ensure sparse labeling) reporter line were imaged from day 1 (30 h post aggregation) to day 2 (43.5 h post aggregation). Imaging was caried out at 26°C directly in low binding 96-well plate (Thermo, Cat#:174925 Nunclon Sphera) with ACQUIFER Imaging Machine (ACQUIFER Imaging GmbH, Heidelberg, Germany) (Pandey et al., 2019). For each well, a set of 30-50 z-slices (9 µm step size) was acquired in the bright field (50% LED intensity, 40 ms exposure time) and 470 nm fluorescence (100 % LED excitation source, FITC channel, 50 ms exposure time) channels. Imaging was performed with 30 min intervals.

Images were processed with Fiji (Schindelin et al., 2012). Cell tracking was performed using Manual tracking with TrackMate plugin in Fiji (Ershov et al., 2022).

### Transcriptome analysis

RNA was isolated from 1 dpf, 2 dpf, 3 dpf and 4 dpf medaka embryos as indicated in Figure 7a. Material from 6 embryos per condition was pooled. For embryos 1 and 2 dpf, head region containing optic vesicle and head structure anterior to the optic vesicles and eye cup and tissue anterior to the eye respectively were dissected in ice cold PBS. For stages 3 dpf and 4 dpf, eyes were dissected in ice cold PBS. Considering 3 h delay in organoid development with the respect to embryonic development (Zilova et al., 2021), organoids were collected for RNA isolation 3 h later than the embryonic tissue. For organoids, 20 organoids were pooled per condition. Samples were prepared in three replicates. Samples were rinsed with ice cold PBS and homogenized in 200 μl of Trizole and RNA was isolated using Direct-zol RNA Microprep (Zymoresearch, Cat#:R2060) according to the manufacture protocol. Library preparation was performed with the NEBNext Ultra II RNA Library Prep Kit for Illumina with NEBNext Poly A Selection Kit and NEBNext Multiplex Oligos with UMI. Input RNA was quantified with the Qubit BR RNA assay, and qualified on the Agilent Tapestation (RNA Assay), libraries were quantified with the Qubit DNA HS Assay, and qualified on the Agilent Tapestion (D1000 Assay). Libraries were sequenced on the Illumina NextSeq 550 Sequencer.

For downstream bioinformatic analysis, UMIs were extracted using UMI-Tools v1.1.2 (Smith et al., 2017). The first 12 bases of each read were trimmed, and reads shorter than 50 bases with an average quality lower than 20 were discarded using Trimmomatic v0.39 (Bolger et al., 2014). Processed reads were mapped to the Japanese medaka HdrR reference genome ASM223467v1 using Bowtie 2 v2.4.4 (Langmead and Salzberg, 2012), and deduplication of the mapped reads was performed with UMI-Tools v1.1.2 (Smith et al., 2017). Gene counts were extracted with featureCounts v2.0.3 (Liao et al., 2014) using the Ensembl version 113 GTF as a reference. The subsequent analysis was performed using R v.4.4.1 (https://www.r-project.org/). Genes with fewer than 10 counts across samples were excluded, and DESeq2 (Love et al., 2014) was used for normalization and variance stabilizing transformation of the gene counts. The transformed counts were then used for principal component analysis and heatmap visualization of lens-specific gene expression.

### Generation of *FoxE3::GFP* reporter line

1.5 kb region upstream of *Foxe3* (ENSORLG00000017669) ATG was PCR amplified from medaka fish genomic DNA with Q5 High Fidelity DNA Polymerase (NEB, Cat#:M0491) using following primers: KpnI-FoxE3 (forward primer): 5’-TAAGGTACCTGGAATCAAGCAGAAATAATCCACCAAAAGCA and FoxE3_R1 (reverse primer): 5’-GGGGTACCGTTGCACATGCCAGAGAAGTAG, sub-cloned into pJet1.2/blunt cloning vector (Thermo Fisher Scientific), released via restriction digest with XhoI (NEB, Cat#:R0146) and NcoI (NEB, Cat#:R0193) restriction enzymes. The plasmid ISceI-pBSII KS+ (NovoPro, Cat#:V009906) (containing GFP sequence inserted via EcoRI and XbaI restriction sites) was digested with XhoI and NcoI, and ligated with the 1.5 kb *Foxe3* upstream regulatory element to generate *I-Sce/FoxE3(1.5kb)::*GFP. Plasmid DNA was isolated with QuiAprep Spin Miniprep Kit (Quiagen, Cat#:27106), sequenced (Eurofins Genomics) and analyzed using Geneious R8.1 (Biomatters). *I-Sce/FoxE3(1.5kb)::*GFP (10 ng/μl) was co-injected with Meganuclease (I-SceI) (NEB, Cat#:R0694L) into the cytoplasm of one cell stage medaka zygotes as previously described (Thermes et al., 2002) to generate *Foxe3::GFP* reporter line. Embryos were raised and maintained at 26°C in 1× Embryo Rearing Medium (ERM, 17 mM NaCl, 40 mM KCl, 0.27 mM CaCl_2_, 0.66 mM MgSO_4_, 17 mM HEPES) and screened for ocular GFP expression 1 dpf on a Nikon SMZ18.

### Quantitative analysis of organoid and lens size

Organoid and lens sizes (Figure 2-figure supplement 1, Figure 5d and 5f) were measured as diameter (mean of 4 measurements per organoid) of the organoid on central optical section through organoid (organoid size at day 1) or on maximal projection of multiple z planes of the organoid (lens size at day 2, 4 or 5). Depth of *Rx3::H2B-GFP* expression (Figure 5e) was measured on central optical section through the organoid (mean of 8 measurements per organoid) as a thickness of tissue carrying GFP^+^ cells. Statistical analysis and plots were prepared with GraphPad. Figures were assembled with Affinity Designer 2.

## Supporting information

DataS1

## Lead contact

- Requests for further information and resources should be directed to the lead contact, Lucie Zilova

## Materials Availability Statement

- Plasmids and fish lines generated in this study will be made available on request, but we might require a completed materials transfer agreement if there is potential for commercial application.

## Data and Code Availability Statement

- Bulk RNA sequencing data have been deposited in publicly available repository of Heidelberg University, HeiData and are publicly available under https://doi.org/10.11588/DATA/VEREK9 as of the date of publication.
- Microscopy data reported in this paper will be shared by the lead contact upon request.
- This paper does not report original code.
- Any additional information required to re-analyse the data reported in this paper is available from the lead contact upon request.

## Acknowledgements

The authors thank to E. Hasel de Carvalho and J. Benjaminsen for valuable discussions and comments on the manuscript, T. Kellner and B. Wittbrodt for technical help and maintaining fish stocks. Library preparation and sequencing was performed by CCTP Deep Sequencing Facility of Heidelberg University.

This work was supported by grants of the Excellence Cluster “3D Matter Made to Order” (EXC-2082/1-390761711) funded through the German Excellence Strategy via Deutsche Forschungsgemeinschaft (DFG, German Research Foundation), PostDoc take-off Grant funded through and the Excellence Cluster ‘3D Matter Made to Order’ to L.Z. and J.W and Dr. Moos-Stiftung. For the publication fee, we acknowledge financial support by Heidelberg University.

## Author Contributions Statement

Conceptualization: L.Z., J.W.

Investigation: E.S., M.A.D.T., L.Z., A.P., I.S.

Visualization: E.S., M.A.D.T., I.S., L.Z.

Funding acquisition: J.W., L.Z.

Project administration: J.W. Supervision: L.Z.

Writing - original draft: L.Z., J.W.

## Supplemental information

Video S1-S8.

Video S1. Cells of the outer layer of the organoid acquire retina fate, related to Figure 3.

Video S2. Time laps imaging of multiple organoids from day 1 to day 2 showing the formation of the lens, related to Figure 4.

Video S3. Lens fate is acquired by cells localized in central part of the organoid, related to Figure 3.

Video S4. Arrangement of retinal and lens tissue in day 2 organoid, related to Figure 4.

Video S5. Re-aggregation of dissociated day 1 organoids, related to Figure S4.

Video S6. Tracking of movement of individual lens progenitor cells from the center to the periphery of the organoid, related to Figure 4.

Video S7. Cell shape changes of lens-committed progenitor cells during the process of lens displacement from center to the organoid periphery, related to Figure 4.

Video S8. Movement of retinal progenitor cells during the process of organoid formation from day 1 to day 2.

Data S1. Excel table, related to transcriptome analysis of Figure 7. Normalized and variance stabilized counts labeled with Ensemble IDs.

---

**Figure 2-figure supplement 1.**
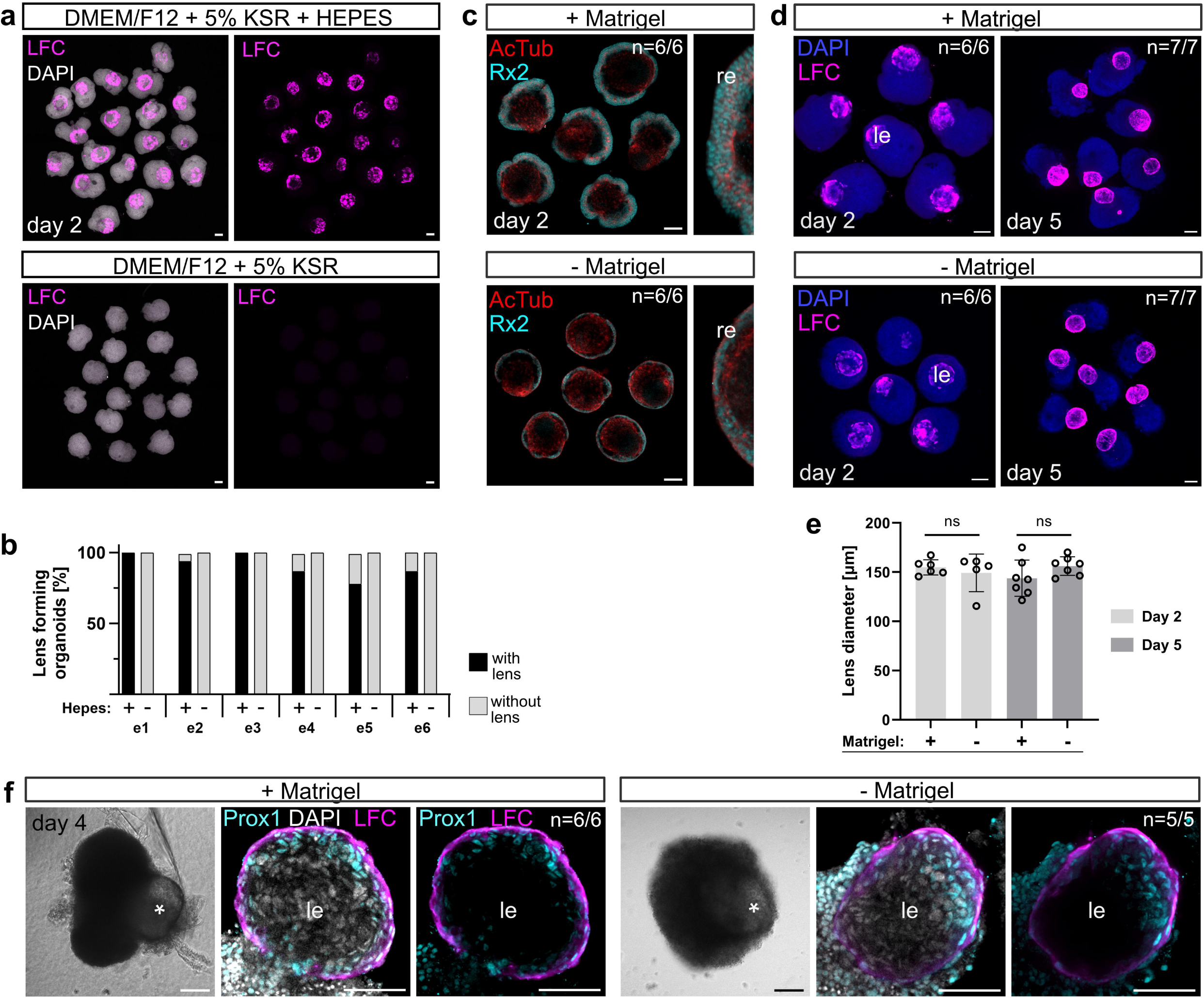
Expression of lens-specific markers is dependent on the presence of HEPES in the media and occurs independently of Matrigel addition, related to Figure 1 and Figure 2. **(a)** Day 2 organoids generated in DMEM/F12 + 5 % KSR media in the presence or absence of 20 mM HEPES labeled with anti-LFC antibody, co-labelled with DAPI nuclear stain. **(b)** Quantification of the efficiency of lens formation measured by the expression of lens marker LFC in day 2 organoids grown in differentiation media with or without HEPES in n=6 independent experiments (e1 – e6). Total number of analyzed organoids: n=63 without HEPES; n=83 with HEPES. **(c-f)** Impact of Matrigel supplementation on organoid organization. **(c)** Day 2 organoids supplemented or not supplemented with Matrigel (2 % final concentration) at day 1 labeled with anti-Rx2 and anti-AcTub (acetylated tubulin) antibody to visualize the structure of retinal neuroepithelium. **(d)** Day 2 and day 5 organoids supplemented or not supplemented with Matrigel (2 % final concentration) at day 1 labeled with anti-LFC antibody (displayed as maximal projections of multiple z planes). **(e)** Quantification of the size of the lens measured as the diameter of LFC^+^ lens sphere in d. Statistical significance was calculated using an unpaired two-tailed t-test. n.s., not significant. **(f)** Day 4 organoids labelled with anti-GFP and anti-LFC antibody, co-labeled with DAPI nuclear stain, represented by single optical section. The effect of Matrigel supplementation was assessed in total of n=49 organoids in 4 independent experiments. n numbers indicate the number of organoids with displayed phenotype within one experiment. LFC, lens fiber cell; re, retina; le, lens. Scale bar in a, c, d, and f: 100 μm, in enlarged lens in f: scale bar 50 μm.

**Figure 3-figure supplement 1:**
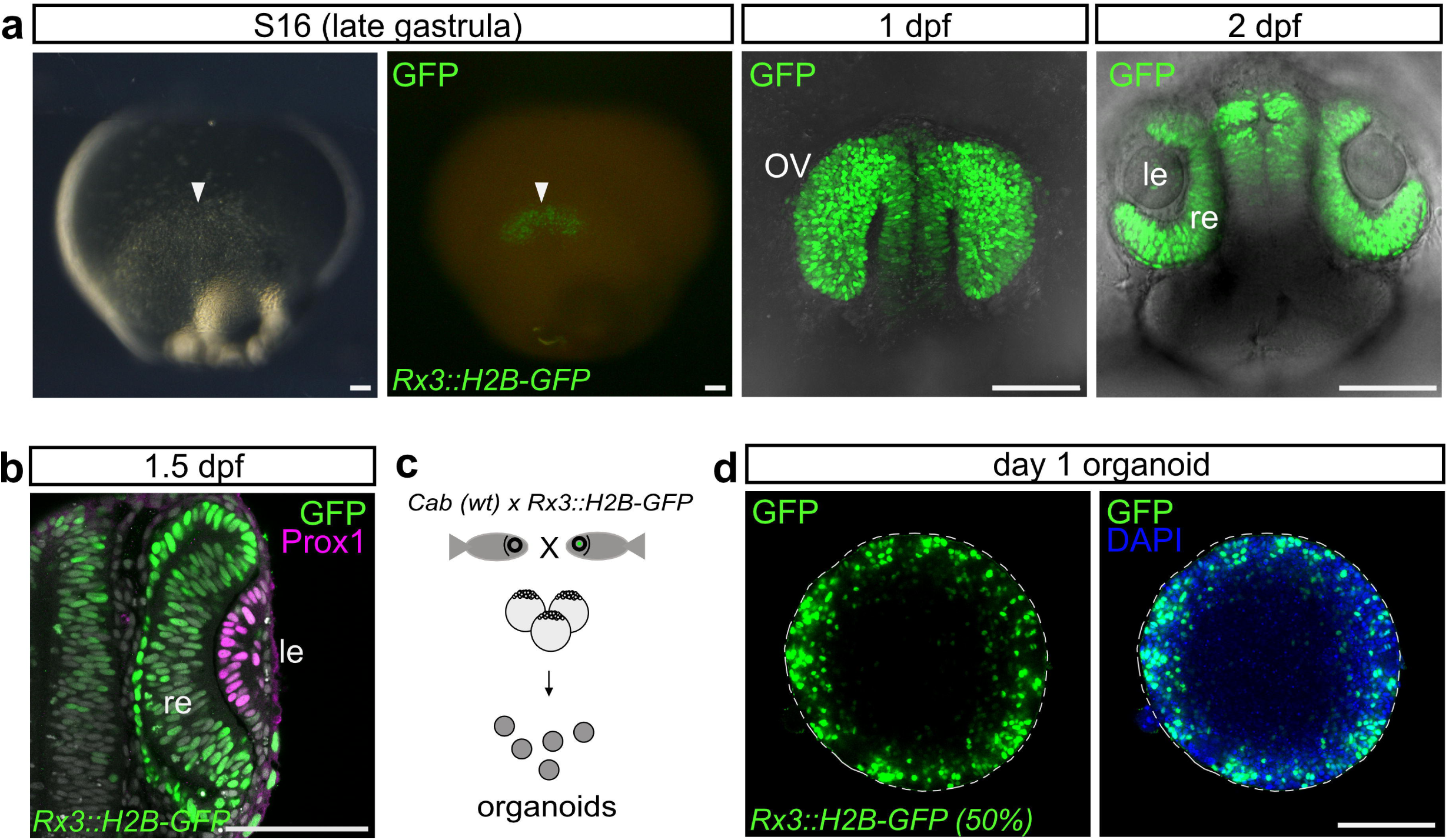
Expression domain of *Rx3::H2B-GFP* in the embryo and organoid, related to Figure 3. **(a)** Expression domain of *Rx3::H2B-GFP* in medaka fish embryo at indicated stages showing the expression in the anterior neural plate - presumptive eye field (S16, late gastrula stage, arrowhead), retina-committed progenitors of the optic vesicle (1 dpf), retina and forebrain (2 dpf). 1 dpf is represented by maximal z-projection. 2 dpf represented by single optical section through a live embryo. **(b)** Optical section through medaka fish embryo 1.5 dpf (S22) immunostained with antibodies against GFP and Prox1 showing lens-specific expression of Prox1 and retina-specific expression of *Rx3::H2B-GFP*. **(c)** Schematic representation of organoid generation from the retina-specific *Rx3::H2B-GFP* reporter line (50 % of cells). **(d)** Single optical section showing peripheral expression of retina specific marker *Rx3::H2B-GFP* in the organoid at day 1 visualized by immunohistochemistry with antibody against GFP; co-stained with DAPI nuclear stain. For *Rx3::H2B-GFP*, the percentage (%) indicates how many cells carry the indicated transgene. Dashed line indicates the outline of the organoid. wt, wild type; le, lens; re, retina; OV, optic vesicle. Images acquired with laser scanning confocal microscope sp8 (Leica). Scale bar 100 μm.

**Figure 3-figure supplement 2.**
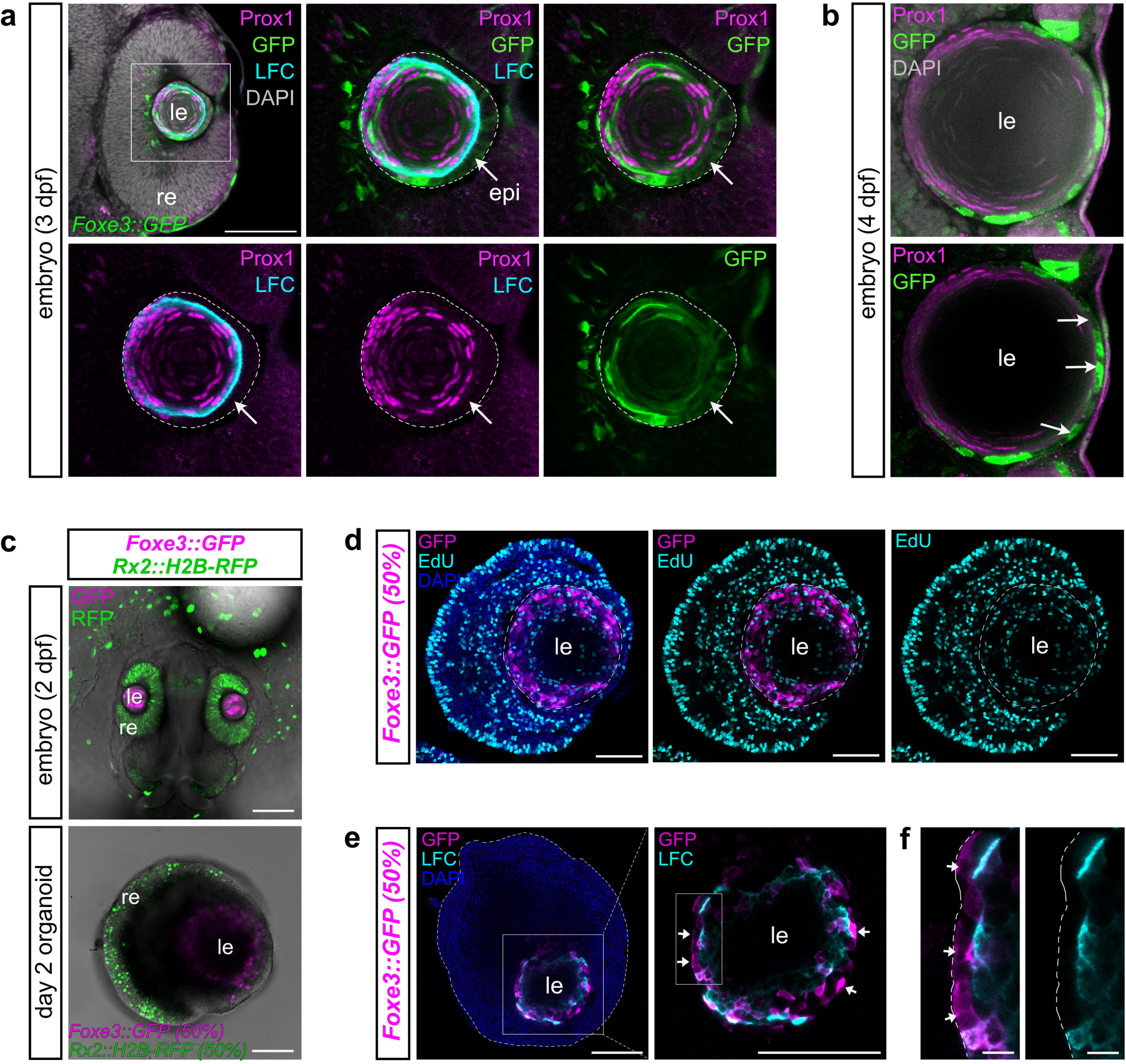
Characterization of *Foxe3::GFP* reporter-expressing cells in embryos and organoids, related to Figure 3. **(a-b)** Expression of *Foxe3::GFP* in medaka embryonic lens. Optical sections of the whole eye and magnification of lens at 3 days post fertilization (a) and lens at 4 days post fertilization (b) showing the expression of *Foxe3*-driven GFP in lens epithelium (arrows) and differentiating lens fiber cells (core of the lens), co-labeled with the antibody against Prox1 and LFC lens fiber cell antibody. Dashed line indicates the outline of the lens in a. **(c)** Expression of *Rx2::H2B-GFP* and *Foxe3::GFP*, in the day 2 embryo and organoid. **(d)** EdU incorporation assay labeling proliferating (S-phase) cells in day 2 *Foxe3::GFP*-derived organoids exposed to EdU for 1 hour, GFP (magenta) and EdU (cyan) were detected by immunolabeling with anti-GFP antibody and EdU detection. **(e)** Distribution of *Foxe3-*driven GFP and lens differentiation marker LFC. Dashed line indicates the outline of the organoid. **(f)** Detail of indicated area in e highlighting *Foxe3::GFP*^+^/LFC^-^ cells, similar to the progenitor-carrying lens epithelium in embryonic lens (arrows). Dashed line indicates the outline of the lens. For *Foxe3::GFP* and *Rx2::H2B-RFP*, the percentage (%) indicates how many cells carry the indicated transgene. le, lens; re, retina; epi, lens epithelium; LFC, lens fiber cell; dpf, days post fertilization. Scale bar 100 μm.

**Figure 4-figure supplement 1.**
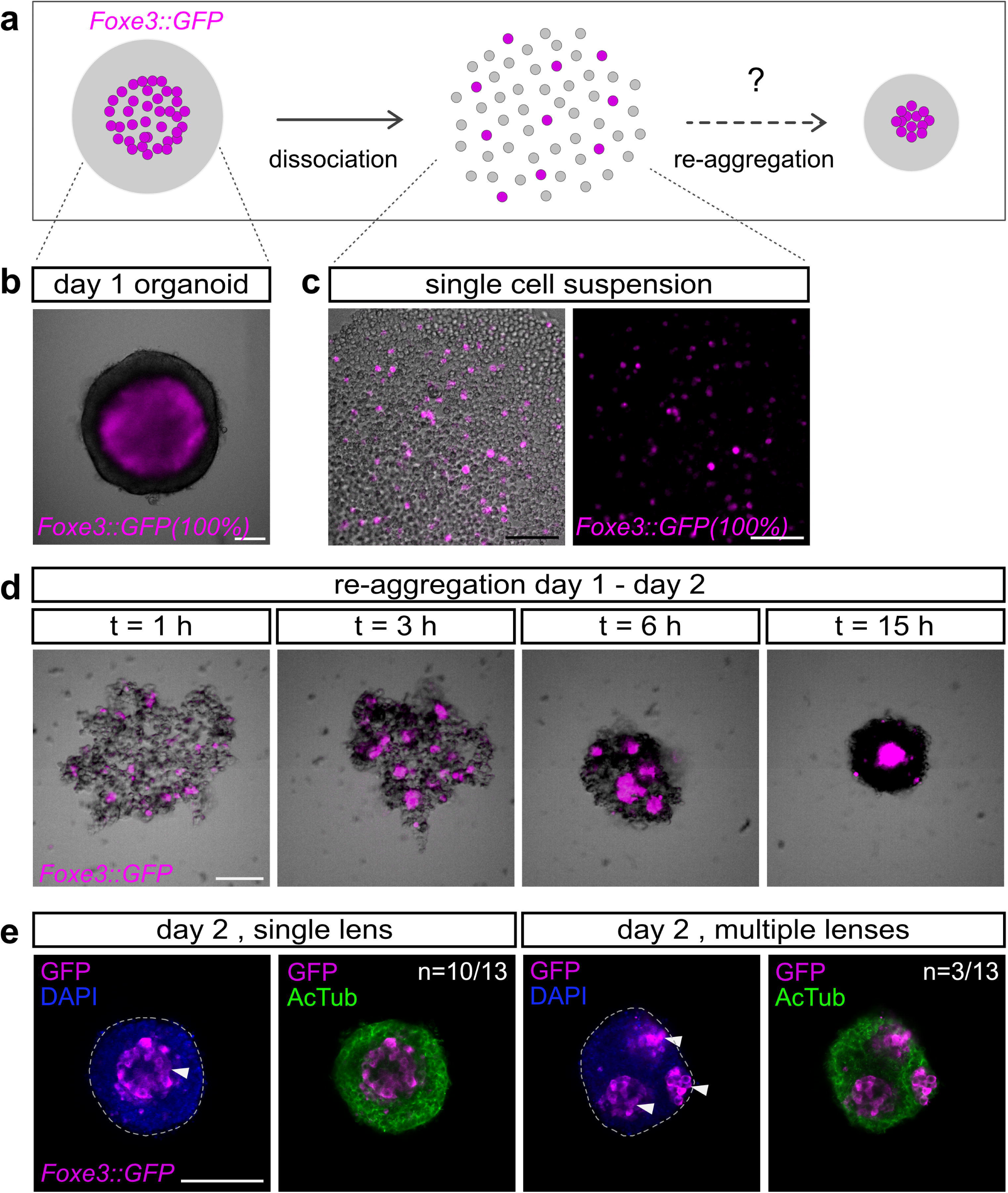
Retinal and lens progenitor cells show differential adhesive properties allowing for cell identity-based cell sorting. **(a)** Schematic representation of dissociation/re-aggregation experiment. Day 1 organoids derived from *Foxe3::GFP* lens reporter line were dissociated into single-cell suspension and left to re-aggregate. **(b)** Representative picture of day 1 *Foxe3::GFP* organoid before dissociation showing lens-specific GFP expression in the central region of the organoid. (c) Single cell suspension generated by trypsin-mediated organoid dissociation. **(d)** Process of re-aggregation showing gradual sorting of *Foxe3::GFP*-expressing lens progenitors. Time (t) indicated in hours post dissociation. **(e)** Representative images of organoids generated by dissociation/re-aggregation process showing distribution of retinal (AcTub^+^) and lens (GFP^+^) cells analyzed by immunohistochemistry. In 3 independent experiments, total number of n=35 organoids were analyzed. n numbers in e indicate the number of organoids with displayed phenotype within one experiment. Scale bar 100 μm.

**Video S1. Cells of the outer layer of the organoid acquire retina fate, related to Figure 3**. Whole volume of day 1 organoid generated form retina-specific *Rx3::H2B-GFP* (50%) reporter line, immunolabeled with anti-GFP antibody, co-stained with DAPI nuclear stain. The percentage (%) indicates the proportion of cells within the organoid carrying indicated reporter. The volume was acquired with 1 µm z step size with confocal microscope Sp8. Scale bar 100 μm.

**Video S2. Time laps imaging of multiple organoids from day 1 to day 2 showing the formation of the lens, related to Figure 4**. Overlay of brightfield and epifluorescence images of organoids generated from *Foxe3::GFP* (50%) reporter line imaged with the 30 min intervals over 16 h from day 1 (24h post aggregation) to day 2 (40h post aggregation). The percentage (%) indicates the proportion of cells within the organoid carrying indicated reporter. Acquired with ACQUIFER Imaging Machine. Scale bar 100 μm.

**Video S3. Lens fate is acquired by cells localized in central part of the organoid, related to Figure 3**. Whole volume of day 1 organoid generated form retina-specific *Foxe3::GFP* (50%) reporter line, immunolabeled with anti-GFP antibody, co-stained with DAPI nuclear stain. The percentage (%) indicates the proportion of cells within the organoid carrying indicated reporter. The volume was acquired with 1 µm z step size with confocal microscope Sp8. Scale bar 100 μm.

**Video S4. Arrangement of retinal and lens tissue in day 2 organoid, related to Figure 3**. Rendering of day 2 organoid generated by mixing of cells from *Rx2::H2B-RFP* (50%) (green) and *Foxe3::GFP* (50%) (magenta) reporter lines immunolabelled with anti-GFP and RFP antibody, co-stained with DAPI nuclear stain (cyan). The percentage (%) indicates the proportion of cells within the organoid carrying indicated reporter. Imaged with Sp8 confocal microscope. Scale bar 100 μm.

**Video S5. Re-aggregation of dissociated day 1 organoids, related to Figure 4**. Reaggregation of cells generated by dissociation of *Foxe3::GFP* (100%) lens reporter-derived day 1 organoids showing sorting of lens/retinal cell populations imaged with the 1 h intervals over 15 h from day 1 to day 2. Acquired with ACQUIFER Imaging Machine. Scale bar 100 μm.

**Video S6. Tracking of movement of individual lens progenitor cells from the center to the periphery of the organoid, related to Figure 4**. Organoids derived from *Foxe3::GFP* lens reporter line (10 % of reporter cells with 90 % of wild-type cells) imaged with the 30 min intervals over 10 h from day 1 to day 2. Acquired with ACQUIFER Imaging Machine. Scale bar 100 μm.

**Video S7. Cell shape changes of lens-committed progenitor cells during the process of lens displacement from center to the organoid periphery, related to Figure 4**. Organoids derived from *Foxe3::GFP* lens reporter line (10 % of reporter cells with 90 % of wild-type cells) imaged with the 30 min intervals over 13 h from day 1 to day 2. Acquired with ACQUIFER Imaging Machine. Scale bar 100 μm; in detail: scale bar 25 μm.

**Video S8. Movement or retinal progenitor cells during the process of organoid formation from day 1 to day 2.** Organoids derived from *Rx3::H2B-GFP* retinal reporter line (2% of reporter cells with 98 % of wild-type cells) imaged with the 30 min intervals over 13.5 h from day 1 to day 2. Acquired with ACQUIFER Imaging Machine. Scale bar 100 μm.

